# aPKC-ζ III promotes trophoblast fusion by altering Par-3 interactions with Hippo Signaling Kinase LATS1

**DOI:** 10.64898/2025.12.05.692666

**Authors:** Sumaiyah Z. Shaha, Wendy K. Duan, Juan Garcia Rivas, Ivan K. Domingo, Meghan Riddell

**Affiliations:** Department of Physiology, University of Alberta, Edmonton AB, Canada; Department of Obstetrics and Gynecology, University of Alberta, Edmonton AB, Canada

## Abstract

The first trimester of pregnancy is a critical developmental period for the placenta. In humans, the maternal-facing exchange surface is formed by a single giant multinucleate syncytium: the syncytiotrophoblast (ST). The ST arises from villous lineage commitment of trophoblast stem cells (TSC) and the differentiation and fusion of progenitor cytotrophoblasts (pCT) to form the multinucleate syncytium. The Hippo signaling co-transcription factor YAP1 promotes pCT maintenance and TSC stemness, however, how Hippo signaling is regulated remains unknown. We have identified a novel *PRKCZ* encoded aPKC isoform, aPKC-ζ III, that is highly expressed in pCT and ST. Here we establish that aPKC-ζ III promotes pCT fusion by regulating Hippo signaling. Specifically, aPKC-ζ III outcompetes the Hippo kinase LATS1 for scaffolding protein Par-3 binding, resulting in YAP1 inactivation and pCT fusion. Our findings identify a key modulator of Hippo signaling in human trophoblasts that is critical for first trimester ST differentiation.

## Introduction

Formation of the placenta is critical for the establishment and progression of pregnancy. It is a fetally derived organ that is responsible for facilitating nutrient transport, gas exchange, and hormone secretion amongst other critical functions. Trophoblasts are placental-specific epithelial cells. Trophoblast stem cells (TSC) are derived from the trophectoderm (TE) of the blastocyst; thus, this represents the first committed cell lineage during development. In humans, the syncytiotrophoblast (ST) is the terminally differentiated cell of the villous trophoblast lineage. It is a giant multinucleate cell that spans the surface of the maternal-facing exchange surface of the placenta. This single giant cell facilitates the transfer of gases and nutrients while secreting pregnancy specific hormones to promote placental development and maternal adaptation to pregnancy.^1,2^ The ST is post-mitotic and relies on the differentiation of underlying progenitor cytotrophoblasts (pCTs) for its expansion and maintenance.^3^ The differentiation of pCT into ST is a complex multistep process that culminates in cell-cell fusion, and upregulation of genes specific for ST function and homeostasis.^4,5^ In common pregnancy complications such as intrauterine growth restriction and preeclampsia, there are defects in pCT to ST fusion that are thought to arise within early placental formation.^6,7^ Thus, understanding pCT to ST differentiation in the first trimester is critical.

The Hippo signaling pathway is involved in regulating organ growth, cell proliferation, and differentiation in many cellular contexts.^8,9^ Hippo signaling is regulated by numerous signals such as cell contact, mechanical cues, stress, and cell polarity.^9^ When Hippo signaling is active, mammalian Ste20-like kinases 1/2 (MST1/2) phosphorylate and activate large tumor suppressor 1/2 (LATS1/2), which in turn phosphorylates co-transcription factors Yes-associated protein 1 (YAP) and transcriptional coactivation for PDZ-binding motif (TAZ), resulting in YAP/TAZ cytoplasmic retention and/or ubiquitination and degradation.^10,11^ When Hippo signaling is disrupted, non-phosphorylated YAP/TAZ (active) translocate to the nucleus and interact with DNA-binding transcriptional enhanced associate domains 1-4 (TEAD1-4) to influence transcription.^12,13^ In human trophoblasts, Hippo signaling has been established as a critical regulator of pCT maintenance.^14,15^ YAP is highly expressed within the nucleus in the pCT population resulting in TEAD4 activity. This leads to the transcription of genes promoting trophoblast stemness and repression of genes necessary for cell fusion and ST differentiation.^14,15^ While it is understood that Hippo signaling is critical for pCT maintenance, what regulates Hippo signaling in trophoblasts has not yet been examined.

During human TE segregation from the inner cell mass (ICM), the outer cells become polarized and YAP1 localizes to the nucleus.^16^ This initiation and maintenance of cell polarity is governed by the activity of an apical-basal polarity regulator, atypical protein kinase C (aPKC). When TE aPKC expression or activity is disrupted YAP1 localization is altered.^16^ Therefore, in the pre-TSC TE, Hippo signaling and cell polarity are linked. Cell polarity regulatory complexes are well known to play important roles in epithelial cell maintenance and differentiation, however, studies examining polarity regulators in the human placenta are limited.^17,18^ The Par complex is an evolutionarily conserved polarity regulating complex that consists of scaffolding proteins partitioning defective-3 (Par-3) and partitioning defective-6 (Par-6), and aPKC isoforms. There are two main isoforms of aPKC in humans: aPKC- and aPKC-ζ encoded by *PRKCI* and *PRKCZ,* respectively. APKCs are spatio-temporally regulated as their full kinase activation is dependent upon protein-protein interactions.^19^ Murine models have revealed that *Prkci* knockout (KO) is embryonic lethal by day 9 due to defects in placental development, but *Prkcz* KO are grossly normal with impairments in NF-κB signaling.^20–22^ Par complex members have also been implicated in human TSC and pCT to ST differentiation.

Sivasubramaniyam *et al.* established that Par-6 negatively regulates trophoblast fusion.^23^ Among some of the first studies performed using human TSC, aPKC- was shown to promote TSC to ST differentiation.^22^ Adding complexity to the canonical Par complex, we recently identified that the human placenta expresses three isoforms of aPKC: aPKC- , aPKC-ζ, and *PRKCZ* encoded aPKC-ζ III.^24^ APKC-ζ III has an N-terminal truncation that results in a loss of the Phox and Bem1 (PB1) domain, but retained expression of the kinase domain, the pseudosubstrate inhibitory region, and the PDZ binding motifs necessary for interaction with Par-3.^24,25^ The PB1 domain of aPKCs are necessary for interaction with Par-6 via PB1-PB1 mediated interactions. PB1 heterodimerization between aPKC and Par-6 allows for full activation of kinase activity by removal of the pseudosubstrate region from the kinase domain and coupling the activity to the plasma membrane.^19,26^ Thus, it is unclear if aPKC-ζ ΙΙΙ is capable of full catalytic activity.^17^ Presently, the function of aPKC-ζ III in trophoblasts is unknown.

Here we show that aPKC-ζ III promotes pCT fusion in first trimester trophoblasts. We identify that aPKC-ζ III interacts with Par-3 to maintain activation of Hippo signaling kinase LATS1 and inactivity of the co-transcription factor YAP1 to promote pCT fusion.

## Results

### *PRKCZ* is upregulated in the villous trophoblast lineage

Our previous work assessed aPKC- and aPKC-ζ protein and mRNA expression in first trimester and term human placentas, and *in vitro* differentiated ST from primary isolated pCT.^24^ However, single cell RNA-sequencing (scRNA-seq) and single nuclei RNA-sequencing (snRNA-seq) of early human placenta and human trophoblast organoids have revealed that multiple transcriptionally distinct pCT and ST states exist in the villous lineage.^27–32^ These include a fusion competent pCT population identified by the high expression of the trophoblast fusogens syncytin-1 and syncytin-2 (encoded by *ERVW-1* and *ERVFRD-1*, respectively) which are necessary for pCT fusion.^3,33–35^ To understand if Par complex members are expressed in this critical state, we utilized the snRNA-seq data from first trimester placentas previously presented by Wang *et al*.^29^ The dataset was visualized using uniform maniform approximation and projection (UMAP) dimensional reduction analysis. A total of 11 clusters were identified from 45,697 nuclei (Figure 1A). Cluster identification based on cell identify was done by analyzing marker gene expression for placental cell and trophoblast subtypes (Figure S1A-E). ^30,36^ Our analyses revealed four different cytotrophoblast states: Bipotential pCTs (*BCAM*^+^, *ITGA6*^+^, *GATA3^+^*), proliferative pCTs (*MKI67*^+^, *ITGA6*^+^, *GATA3^+^*), pCTs (*ITGA6*^+^, *GATA3*^+^, *MKI67*^-^), and fusion competent pCTs (*ERVW-1*^+^, *ERVFRD-1^high^, GREM2*^+^). We additionally identified two ST subtypes: early ST (*SDC1*^+^, *ERVW-1*^+^, *ERVFRD-1*^low^) and ST (*PAPPA*^+^*,SDC1*^+^).

**Figure 1:**
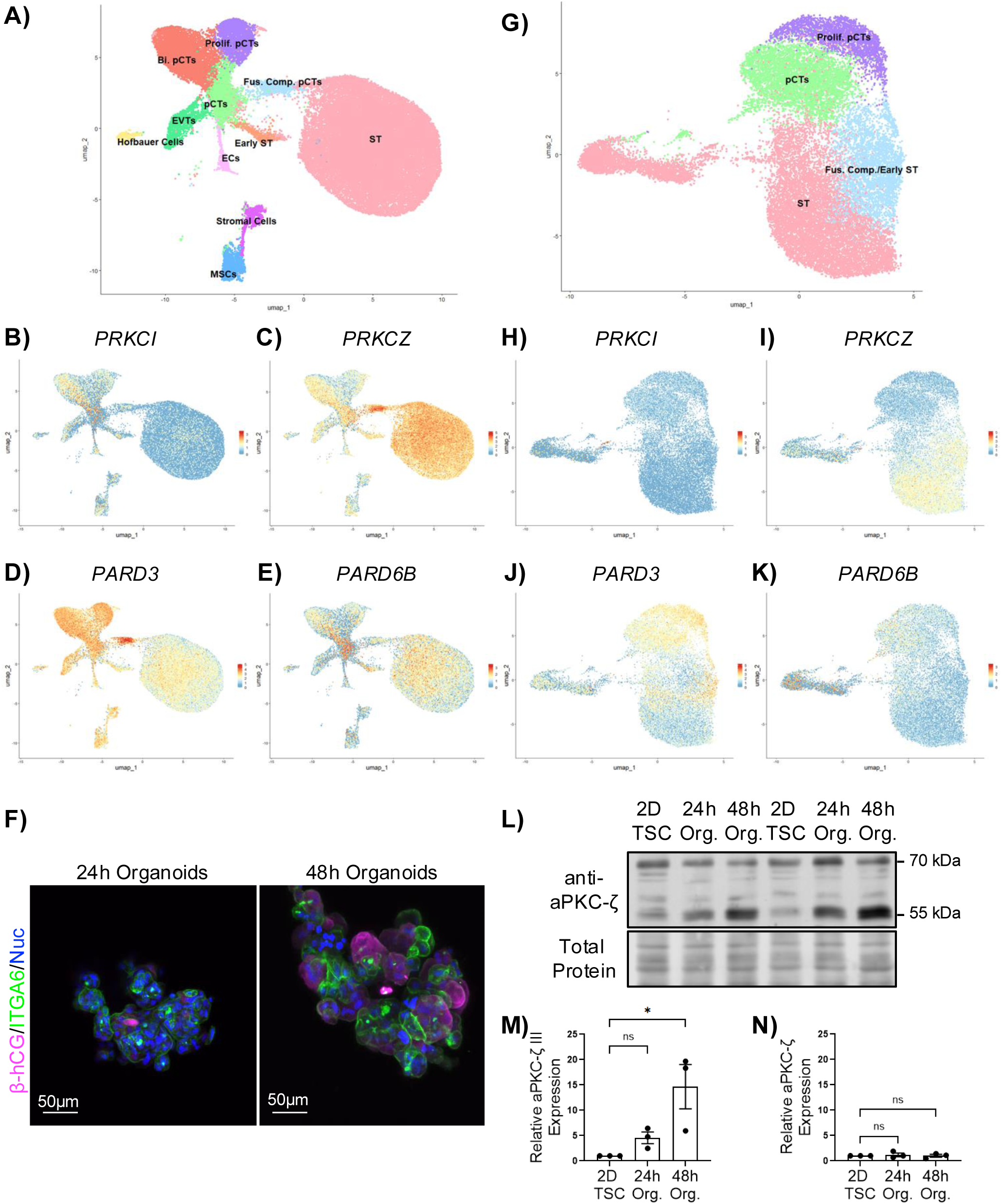
***PRKCZ* encoded aPKC isoforms have increased expression in the villous lineage.** (A) UMAP of nuclei from early first trimester placenta. (B-E) Dimensional reduction plots of (B) *PRKCI*, (C) *PRKCZ*, (D) *PARD3*, and (E) *PARD6B*. F) Representative Z-projection images of 24 and 48 hour ST-organoids stained for ITGA6 (green), β-hCG (magenta), and nuclei (blue). (G) UMAP of nuclei from 48 hour human trophoblast organoids. (H-K) Dimensional reduction plots of (H) *PRKCI*, (I) *PRKCZ*, (J) *PARD3*, and (K) *PARD6B*. (L) Representative western blot of aPKC-ζ and aPKC-ζ III in 2D TSCs, 24 hour and 48 hour organoids. (M and N) Summary data of relative (M) aPKC-ζ III and (N) aPKC-ζ expression; Kruskal-Wallis test with Dunn’s multiple comparisons test, *p<0.05, *n*=3. All graphs show mean +/- S.E.M.

Gene expression analyses revealed *PRKCI* expression in all pCT populations and ST, with the highest density of expression in pCTs, bipotential pCTs and proliferative pCTs, confirming previous work by Bhattacharya *et al*. (Figure 1B).^22^ *PRKCZ* expression was observed in all pCT populations, with the highest density of expression in fusion competent pCTs and ST (Figure 1C).^24^ *PARD3* was expressed in all pCT populations and ST, and like *PRKCZ*, the highest density of expression was observed in the fusion competent pCTs (Figure 1D). *PARD6B* was expressed in all pCT populations, and in ST (Figure 1E). Thus, Par complex members are present in trophoblast villous lineage populations.

To understand if our previously published bioreactor-based trophoblast organoid model^36^ recapitulated first trimester pCT and ST populations, we performed snRNA-seq on organoids derived from human TSC CT27 and CT29 lines after 48 hours of culture, a time point where a mixture of mononucleate pCT populations and multinucleate ST-like cells were present (Figure 1F). We identified four trophoblast clusters from 22,250 nuclei: proliferative CTs (*MKI67^+^ ITGA6*^+^, *ITGA2*^+^), pCTs (*ITGA6*^+^, *GATA3*^+,^ *MKI67^-^*), ST (*ERVW-1*^+^, *ERVFRD-1*^low^, *CGB3*^+^*, GATA3^-^*) and a cluster that had features of both fusion competent and early ST clusters in the analysis of first trimester tissue (Figure 1G, Sup. Fig. 2A-E). Unlike first trimester tissue, no clusters showed a clear fusion competent pCT state with high expression of *GREM2* and low expression of ST marker genes like *SDC1* and β-human chorionic gonadotropin encoding genes (Figure S2).

**Figure 2:**
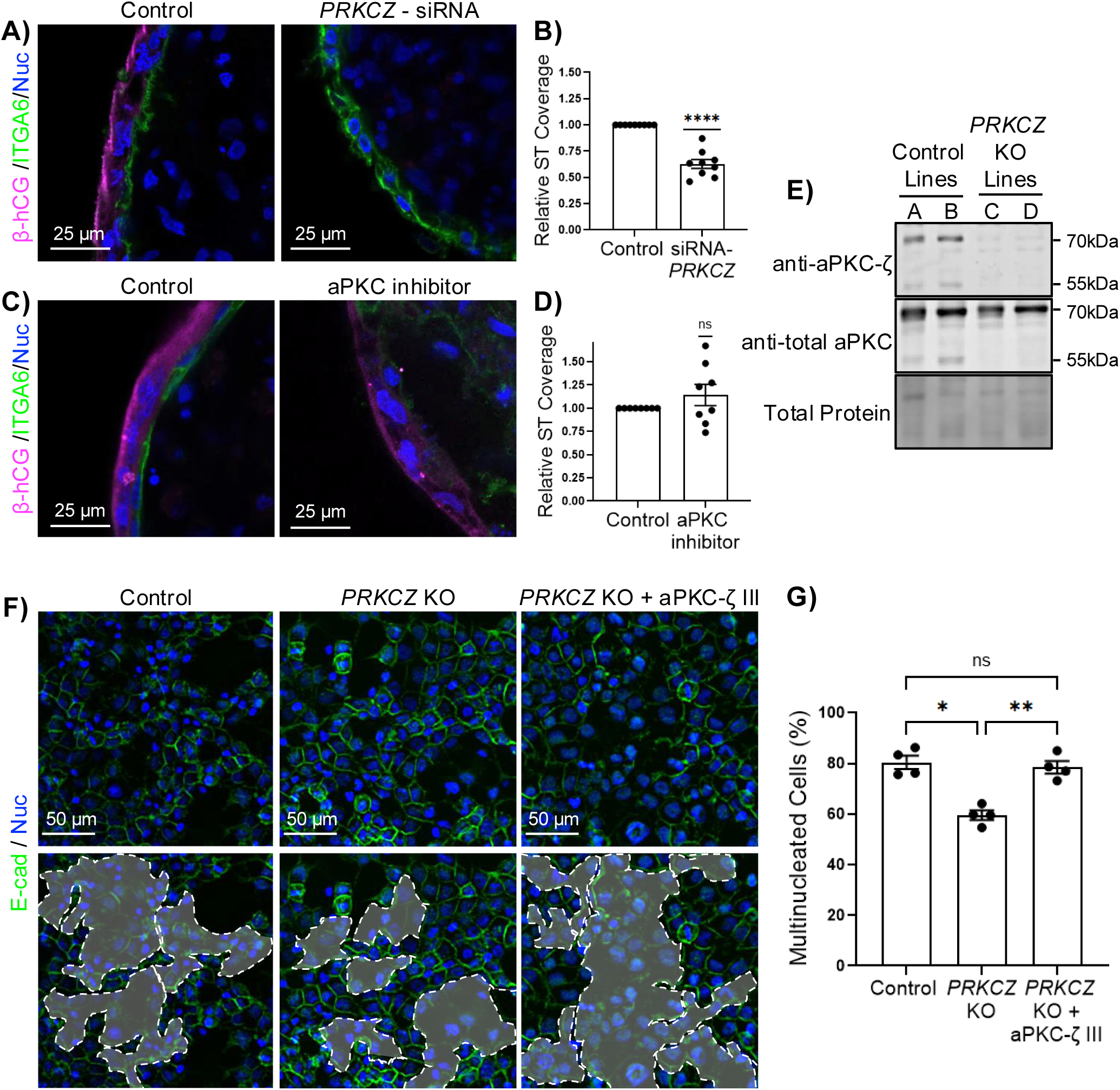
**aPKC-ζ III regulates trophoblast fusion** (A-D) Representative XY-plane images of 9-12 week placenta explants 48 hours post ST-denudation of (A) control and *PRKCZ*-siRNA or (C) control and aPKC inhibitor treated tissue. Summary data of relative ST coverage of (B) *PRKCZ*-siRNA or (D) aPKC inhibitor treated explants; one sample t-test, ****p ≤ 0.0001, n=8-9. (E) Representative western blot of control and aPKC-ζ III KO BeWo cell lines with anti-aPKC-ζ and anti-total aPKC antibodies. (F) Representative images of E-cadherin (green) and nuclei (blue) in control, and *PRKCZ* KO lines +/- aPKC-ζ III rescue; dashed regions in lower panels indicate regions of multinucleated cells. (G) Summary data of percentage of multinucleated cells; one-way ANOVA with Tukey’s multiple comparisons test; *p ≤ 0.05, **p ≤ 0.01; n=4. All graphs show mean +/- S.E.M.

Rather, a large cluster was present with mixed expression of key fusion competent marker genes, like *ERVFRD-*1, which is essential for trophoblast fusion,^35^ and ST markers *SDC1,* and *CGB2* (Figure S2). Together, these data suggest that the organoids recapitulate many signatures of pCT and ST nuclei in intact tissue.

Organoid snRNA-seq data was then used to examine the expression of Par complex components. Gene expression analyses for *PRKCI* revealed expression in pCT and ST populations (Figure 1H). Similar to first trimester tissue, *PRKCZ* was expressed in pCT clusters, with higher expression in the fusion competent/early ST cluster and ST (Figure 1I). *PARD3* and *PARD6B* were variably expressed in all clusters (Figure 1J-K).^30^ Interestingly, despite loss of *Prkcz* having no phenotypic impact on murine placental development,^21^ snRNA-seq expression profiles in both first trimester tissue and organoids suggests *PRKCZ* encoded aPKC isoforms are strongly expressed in villous lineage CTs and are highly expressed in the critical fusion competent state.

We previously observed a non-significant increase in aPKC-ζ III but not aPKC-ζ expression by western blotting and RT-PCR in isolated primary pCTs and *in vitro* differentiated ST from first trimester placenta.^24^ To address if aPKC-ζ III is upregulated during pCT to ST differentiation in human trophoblast organoids, we assessed organoids after 24 and 48 hours of rotational culture to capture a primarily mononucleate pCT population (24 hours) and the progression towards ST-like cell states (48 hours), and compared them to aPKC-ζ/-ζ III levels in undifferentiated TSC (Figure 1F, L-N). Like primary *in vitro* differentiated ST, we observed a consistent and significant 14-fold increase in the ∼55kDa aPKC-ζ III band by western blotting, but not the 70kDa aPKC-ζ band as cells progressed from TSC to 48 hour organoids (Figure 1L-N). Therefore, *PRKCZ* encoded isoforms increase in expression along the villous lineage and display high levels of expression in the critical fusion competent pCT state, suggesting they may play a role in pCT to ST differentiation.

### aPKC-**ζ** III promotes trophoblast fusion

To understand the contribution of *PRKCZ* encoded aPKC-ζ isoforms during trophoblast fusion, we utilized multiple complementary *in vitro* models of pCT fusion and aPKC targeting strategies. We used an *ex vivo* first trimester placental explant ST regeneration model that enriches a fusion competent pCT subpopulation, primary first trimester pCT and BeWo trophoblastic cell lines that fuse and form ST-like multinucleate cells after treatment with cAMP analogs, ^37^ and our trophoblast organoid model.^36^ First trimester placental explants were denuded of ST, and exposed fusion competent pCT were treated with *PRKCZ*-targeting or control siRNA. Treatment with *PRKCZ*-targeting siRNA significantly reduced both aPKC-ζ and aPKC-ζ III expression in explant lysates (Figure S3A-C), and impaired ST regeneration (Figure 2 A,B). Interestingly, when denuded explants were treated with aPKC inhibitor, there was no significant effect on ST regeneration (Figure 2 C,D), suggesting the effects observed with *PRKCZ* knockdown (KD) are independent of kinase activity. APKC inhibitor treatment also had no effect on pCT fusion in primary isolated first trimester trophoblasts induced to fuse for 72 hours using 8-Br-cAMP in 2D culture (Sup. Fig.4).^30^

*PRKCZ* KD was also performed in CT29 TSC and trophoblast organoids were subsequently formed. *PRKCZ*-siRNA treatment resulted in significant reductions in organoid aPKC-ζ and −ζ III (Figure S5A-C) expression, but pCT fusion was unchanged (Figure S5D,E); though *PRKCZ* KD resulted in reduced expression of the ST marker gene *CGB* and trending decrease *GCM1* expression (Figure S5F,G). Suggesting in this model, disruption of *PRKCZ* encoded proteins does not impact pCT fusion.

We used CRISPR-Cas9 technology to target *PRKCZ* in the BeWo trophoblastic cell line (Figure S6).^38,39^ Western blotting revealed the loss of the 55kDa band in two lines compared to two control lines using both an aPKC-ζ specific and total aPKC antibody, with loss of the 70kDa band observed with the aPKC-ζ specific antibody alone (Figure 2E). To determine if *PRKCZ* KO also leads to a decrease in fusion as observed in siRNA treated explants, control and *PRKCZ* KO cells were treated with 8-Br-cAMP to induce fusion. The proportion of multinucleate cells was significantly reduced in *PRKCZ* KO compared to control cells (Figure 2 F,G). The sum of the knockdown, knockout, and aPKC inhibitor data across multiple models suggest that *PRKCZ* encoded isoforms play a kinase activity independent role in regulating pCT fusion.

APKC-ζ III is predicted to have minimal kinase activity due to the absence of the PB1 domain, therefore we hypothesized it may be responsible for the altered pCT fusion we observed in our models. Plasmid-mediated reintroduction of aPKC-ζ III into the *PRKCZ* KO cells rescued fusion to control levels (Figure 2 F,G), suggesting aPKC-ζ III is the *PRKCZ* isoform regulating trophoblast fusion and identifying a function for this newly identified aPKC family member.

### Par-3 and aPKC-ζ III form stable interactions and promote trophoblast fusion

To understand how aPKC-ζ III may be regulating pCT fusion, we examined if the canonical aPKC binding partner Par-3 interacts with aPKC-ζ III and influences pCT fusion, since aPKC-ζ III retains known Par-3 interacting domains.^16^ Par-3 localization has not been reported in first trimester placenta. Thus, to determine if Par-3 and aPKC-ζ isoforms localize to the same compartments, we stained placental tissue from mid to late first trimester with anti-Par-3 antibody. Par-3 signal was localized to pCT E-cadherin junctions, as well as pCT cytoplasm, and inconsistent signal was also observed in ST and stromal cell populations (Figure 3A). The predominant localization of aPKC-ζ III is predicted to be cytoplasmic due to the lack of the PB1 domain required to interact with membrane localized Par-6.^26^ To confirm this, aPKC-ζ III-EGFP was transfected into cells, where EGFP signal was restricted to the cytoplasm as predicted (Figure S7). We had previously reported a strong cytoplasmic anti-aPKC-ζ signal in villous trophoblasts^24^ and, together, these data support that in first trimester pCT, a substantial cytoplasmic pool of aPKC-ζ III and Par-3 exist. To determine if Par-3 and aPKC-ζ III interact, immunoprecipitations were performed. Par-3-EGFP and aPKC- -FLAG (positive control) or aPKC-ζ III-FLAG were expressed in cells and Par-3-EGFP was immunoprecipitated (Figure 3B). Western blot analysis of immunoprecipitated protein revealed aPKC-ζ III can form stable interactions with Par-3 (Figure 3B). As expected, canonical binding between Par-3 and aPKC- was also observed (Figure 3B). To determine if Par-3 is involved in trophoblast fusion in the same pathway as aPKC-ζ III, *PARD3* KD was performed in control and *PRKCZ* KO BeWo cells. Two different *PARD3* targeting siRNAs significantly reduced Par-3 expression in BeWo (Figure S8). *PARD3* KD modestly reduced fusion in control lines, but not in *PRKCZ* KO lines (Figure 3D,E), though there was an additive effect of both *PARD3* KD and *PRKCZ* KO. This suggests Par-3 participates in multiple pathways that modulate trophoblast fusion, and aPKC-ζ III acts downstream of Par-3. Together, our data confirms that Par-3 and aPKC-ζ III interact and are involved in trophoblast fusion.

**Figure 3:**
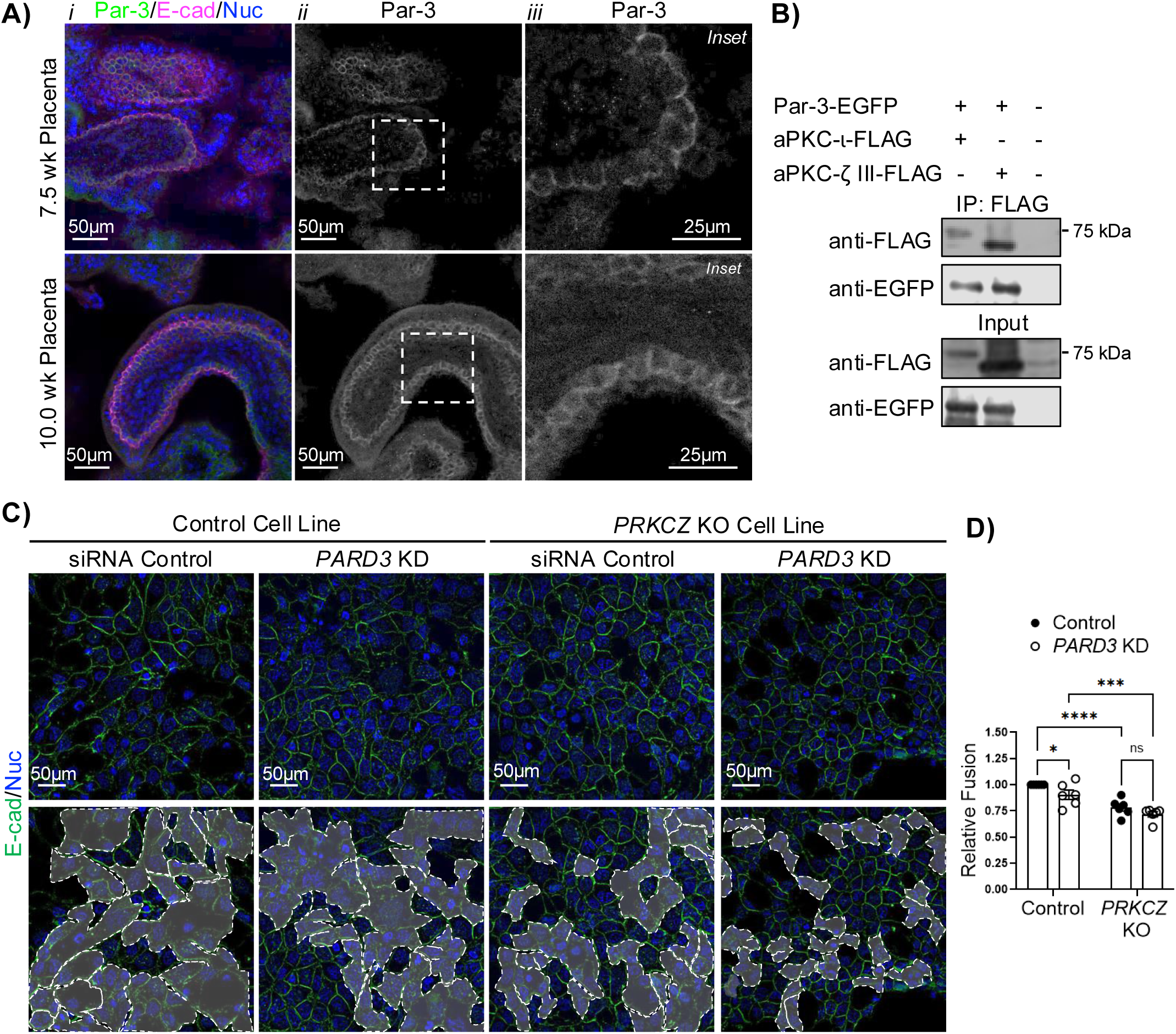
**Par-3 interacts with aPKC-ζ III and promotes trophoblast fusion.** (A) Representative XY-plane images of first trimester placenta tissue stained (*i*) Par-3 (green), E-cad (E-cadherin; magenta) and nuclei (blue); (*ii*) isolated Par-3 signal; (*iii*) Higher magnification inset of isolated Par-3 signal. (B) Western blotting of FLAG immunoprecipitation of Par-3-EGFP and aPKC- -FLAG or aPKC-ζ III-FLAG. (C) Representative images of E-cadherin (green) and nuclei (blue) in control and *PRKCZ* KO BeWo lines treated with control or *PARD3*-targetting siRNA; dashed regions in lower panels indicate regions of multinucleated cells. (D) Summary data of relative fusion; two-way ANOVA with uncorrected Fisher’s LSD; *p ≤ 0.05, ***p ≤ 0.001, ****p ≤ 0.0001; n=6. All graphs show mean +/- S.E.M.

**Figure 4:**
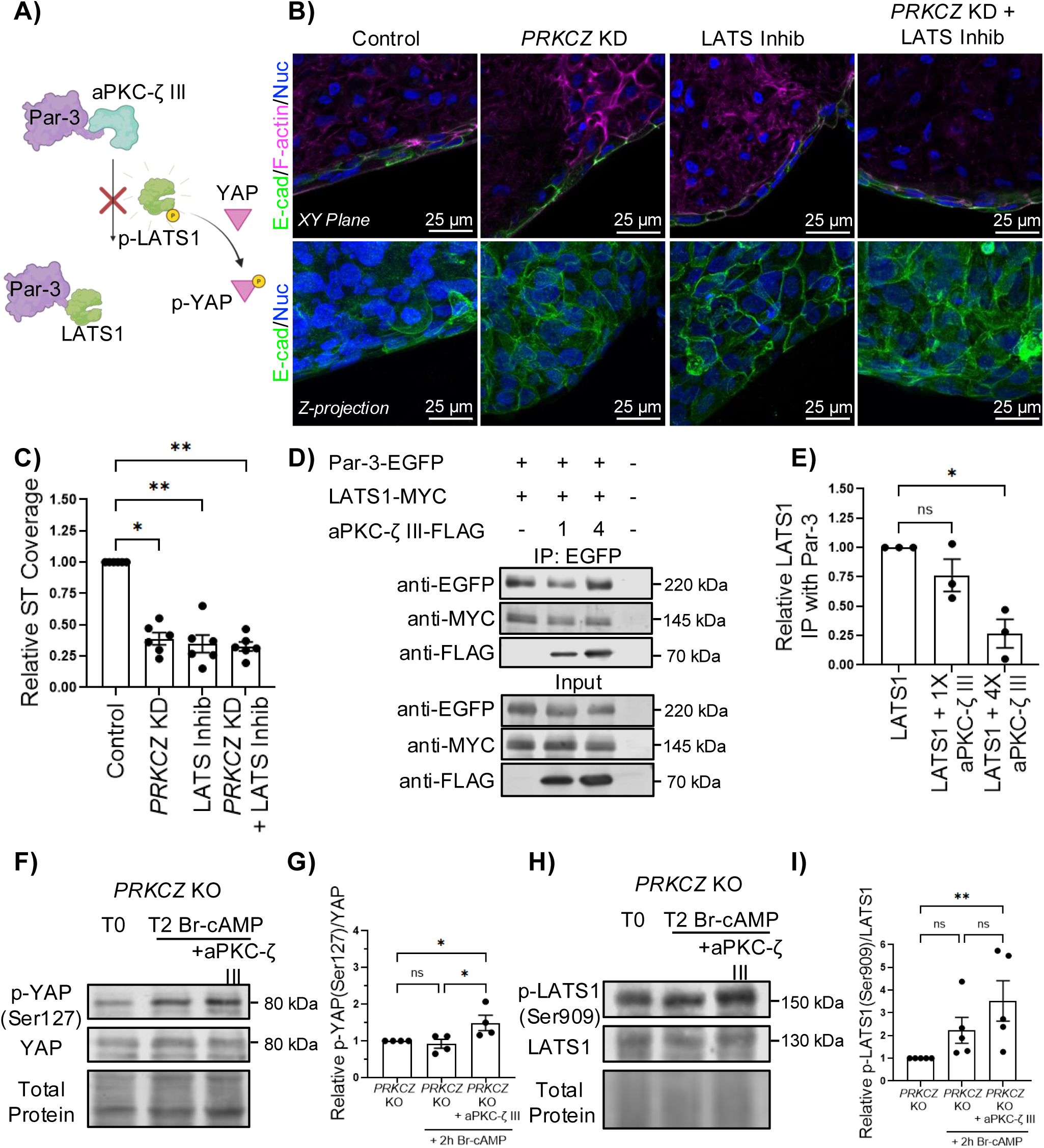
**Hippo signaling is altered by aPKC-ζ III in trophoblasts** (A) Representative XY-plane (top) and Z-projection (bottom) images of 9-12 week placenta explants 48 hours post ST-denudation and treated with *PRKCZ*-siRNA, LATS inhibitor, or a both stained for E-cad (E-cadherin; green), phalloidin (magenta), and nuclei (blue). (B) Summary data of relative ST coverage; Krusal-Wallis test with Dunn’s multiple comparisons; *p ≤ 0.05, **p ≤ 0.01; n=6. (C) Graphical depiction of hypothesized interactions between aPKC-ζ III and Par-3 resulting in phosphorylation of YAP. (D) Western blotting of EGFP immunoprecipitation of Par-3-EGFP with LATS1-Myc +/-aPKC-ζ III-FLAG. (E) Summary data of relative LATS1 immunoprecipitated with Par-3; Kruskal-Wallis test with Dunn’s multiple comparisons test; *p ≤ 0.05; n=3. (F and H) Representative western blot of (F) p-YAP (phospho-Ser127) and total YAP signal or (H) p-LATS (phospho-Ser909) and total LATS from aPKC-ζ III KO cells at T=0 (Control) or T=2 after Br-cAMP +/- aPKC-ζ III reintroduction. (G and I) Summary data of relative (G) p-YAP/total YAP and II) p-LATS1/total LATS1; Kruskal-Wallis test with Dunn’s multiple comparisons test; *p ≤ 0.05, **p ≤ 0.01; n=4-5. All graphs show mean +/- S.E.M.

### aPKC-ζ III modulates the Hippo signaling pathway to promote trophoblast fusion

Cytoplasmic Par-3 has been identified as a modulator of the Hippo signaling pathway by binding Hippo signaling components via its aPKC binding domain.^40^ Specifically, Par-3 promotes the dephosphorylation of LATS1 (inactive) by recruiting protein phosphatase-1 (PP1A), resulting in dephosphorylation of YAP (active) and YAP nuclear translocation.^40,41^ Hippo-YAP signaling has been established as a critical pathway for the maintenance of trophoblast stem state and YAP1 is strongly expressed in both the nuclear and cytoplasmic compartments of pCT of first trimester placenta.^14^ In human TSC, YAP1-TEAD4 complexes promote trophoblast expansion by activating genes associated with trophoblast proliferation but also repress transcription of cell fusion and ST-promoting genes.^14,42^ Therefore, we assessed if *PRKCZ* isoforms are interacting with the Hippo signaling pathway. Though *PRKCZ* KD had no effect on fusion in our organoid model, bulk RNA-sequencing results revealed that differential expression of *CCND1*, a proliferation promoting cell-cycle gene involved in G1-S phase transition, increased in *PRKCZ* KD organoids (Figure S9A). *CCND1* is a direct downstream target gene of YAP1 in the Hippo signaling pathway (Figure S9A).^42,43^ The overall effect of *PRKCZ* KD in organoids was mild with only 16 significant differentially expressed genes observed, precluding Gene Ontology (GO) pathway and Gene Set Enrichment Analysis (GSEA) analyses (Figure S9A). Nuclear and cytoplasmic YAP1 signal was observed in control organoids at multiple time points during organoid maturation (Figure S9B).

Interestingly, pseudotime analyses of trophoblast progenitor differentiation in the organoids using R package Monocle3^44^ predicted multiple pathways leading to ST nuclear states, including some trajectories that do not transit through the pre-fusion/early ST state, but transition directly from a pCT to ST-like state (Figure S10). The imputation of similar differentiation trajectories that skip the pre-fusion pCT state has also been previously reported in scRNA-seq analyses of TSC matrigel organoids.^28^ All together, the combination of these data suggest that Hippo signaling is active in our trophoblast organoid model and may influence pCT differentiation and fusion, but *PRKCZ* isoforms do not influence fusion and are unlikely to modulate Hippo signaling in this model.

As previously mentioned, the aPKC binding region of Par-3 is responsible for increasing the association of LATS1 and YAP1 with PP1A, resulting in decreased kinase activity of LATS1 and subsequent activation of YAP1.^40^ Full length aPKCs bind the Par-3 aPKC-binding region or PDZ2 domains via kinase and PBM domain interactions, both of which are conserved in aPKC-ζ III.^25,45,46^ Thus, we propose a model where aPKC-ζ III binding to Par-3 during pCT fusion and ST formation results in increased LATS1 activity and subsequent YAP phosphorylation and inactivation due to aPKC-ζ III out competing Hippo components for Par-3 binding (Figure 4A).

We utilized our first trimester ST regeneration explant model to confirm that LATS inhibition (YAP1 activation) impaired fusion.^15^ We also sought to determine the effect of combined treatment of LATS inhibitor and *PRKCZ* KD on pCT fusion, because in this model *PRKCZ* KD had the strongest outcome on fusion. TDI-011536 is a derivative of the TRULI LATS inhibitor that blocks LATS kinase activity at nanomolar concentrations, thereby decreasing YAP1 phosphorylation and allowing for subsequent YAP1 nuclear localization.^47^ Therefore, this inhibitor directly targets the regulatory point where aPKC-ζ III is proposed to influence the Hippo signaling pathway. ST denuded placental explants were treated with *PRKCZ-*targeting siRNA, LATS inhibitor, or both for 48 hours (Figure 4B,C). Control explants revealed the regeneration of multinucleated cells on the periphery of explants (Figure 4B). In *PRKCZ* KD explants and LATS inhibitor treated explants there was a 61% and 65% decrease in fusion, respectively, compared to controls, and no additive effect was observed with combined siRNA and inhibitor treatment (Figure 4B,C), suggesting aPKC-ζs and LATS are functioning at the same step of pCT fusion.

To directly test this if aPKC-ζ III reduces LATS1 binding to Par-3, co-IPs were performed with exogenously expressed Par-3, LATS1, and aPKC-ζ III (Figure 4D). Addition of 1:1 ratios of LATS1-MYC:aPKC-ζ III-FLAG plasmid with Par-3-EGFP revealed a non-significant decrease in relative LATS1 immunoprecipitated with Par-3 (Figure 4D-E).

When the ratio of LATS1-MYC:aPKC-ζ III-FLAG plasmid was increased to 1:4, there was a 73% decrease in the relative amount of LATS1 immunoprecipitated with Par-3 (Figure 4D-E), showing that aPKC-ζ III can outcompete LATS1 for Par-3 binding as hypothesized.

To confirm that aPKC-ζ III alters the Hippo signaling pathway during CT fusion, we reintroduced aPKC-ζ III into *PRKCZ* KO BeWo cells and examined phosphorylated levels of LATS1 and YAP. Phosphorylation of YAP at residue Serine 127 promotes YAP cytoplasmic sequestration and inactivation.^48^ APKC-ζ III-FLAG was reintroduced into *PRKCZ* KO cells then *PRKCZ* KO cells +/- aPKC-ζ III-FLAG were treated with 8-Br-cAMP to induce fusion.^39^ After two hours, p-Ser127-YAP remained unchanged in *PRKCZ* KO cells treated with 8-Br-cAMP but increased 1.5-fold in *PRKCZ* KO cells when aPKC-ζ III was rescued relative to untreated controls (Figure 4F,G). Similar experiments were performed to analyze LATS1 phosphorylation at Ser909, essential for LATS1 activation and kinase activity.^10^ Treatment with 8-Br-cAMP alone resulted in a non-significant increase of p-LATS1(Ser909) after two hours which increased 3-fold when aPKC-ζ III was reintroduced (Figure 4H-I). These results indicate aPKC-ζ III modulates the Hippo signaling pathway by reducing LATS1 binding to Par-3 and subsequently inactivates YAP1 to promote pCT fusion and ST differentiation in trophoblasts.

## Discussion

The first trimester of pregnancy is a critical period of development for the human placenta involving the establishment and maintenance of pCT and ST populations. ST malformation underlies many pregnancy complications, thus, understanding the molecular mechanisms that contribute to pCT to ST differentiation remains crucial to identifying therapies. Here, snRNA-seq analyses of human placenta and human trophoblast organoids identify that *PRKCZ* expression is highest in the villous lineage, with elevated expression in fusion competent pCTs/early ST and ST. These data highlight that *PRKCZ* isoforms likely play a role in ST formation, specifically at the fusion competency stage. Using multiple models of pCT fusion, we identified that *PRKCZ* isoforms play an important role in this critical process. Fusion in *PRKCZ* KO cells was rescued by the reintroduction of aPKC-ζ III, revealing that aPKC-ζ III is the aPKC-ζ isoform necessary for regulation of pCT fusion. Finally, we identified that aPKC-ζ III reduces LATS1 binding to Par-3, thereby promoting LATS1 activity, YAP1 inactivation, and establishing a pathway via which aPKC-ζ III regulates ST formation. Our work is the first to establish a key regulator of Hippo signaling in trophoblasts, and we have discovered that this previously unidentified aPKC isoform can modulate Hippo signaling machinery to control cell fate.

The expansion of models to study ST differentiation and advancement of RNA sequencing technologies has led to the identification of multiple distinct pCT states during TSC to ST differentiation.^27,29,30,32,37,49,50^ ScRNA-seq has been performed on human placental tissue and organoid models, however, due to the need for single-cell dissociation, the most differentiated villous trophoblast populations captured are mononucleate precursor ST with very few multinucleate ST.^27,28,32,51,52^ SnRNA-seq has only recently been performed in human placental tissue, capturing for the first time the genetic heterogeneity within the multinucleated ST.^29,30^ To the best of our knowledge, we are the first to perform snRNA-seq on TSC-derived organoids, allowing for better resolution and fidelity of ST populations captured *in vitro.* The availability of multiple different models has unveiled the potential to recapitulate different pCT states, each with their own strengths to model transitionary stages during ST differentiation. ^28,30,53^ Our data highlights how molecules may contribute to ST differentiation at specific steps, and that using a combination of models is critical to address the role of individual molecules in villous differentiation. Previous studies identified that aPKC- is important for human TSC to ST differentiation, and we observed the highest density of *PRKCI* expression in bipotential cells in our snRNA-seq analysis of first trimester tissue, whereas *PRKCZ* isoforms and *PARD3* mRNA are upregulated in fusion competent pCTs (Figure 1A-C).^22,29^ This suggests that aPKC-ι may play a regulatory role earlier in the differentiation pathway towards ST, whereas aPKC-ζ isoforms and Par-3 modulate the critical fusion competence regulatory point during differentiation (Figure 1B-C).

Therefore, with the availability of numerous models in the trophoblast field, it will be important for researchers to use a combination of methods to interrogate the full spectra of regulatory pathways that may occur *in vivo* and contribute to trophoblast differentiation.

SnRNA-seq of human first trimester placenta tissue from Wang *et al.* and bioreactor trophoblast organoids revealed that while organoids remain an excellent 3D model to understand complex interactions between cells, not all pCT subtypes are represented in high abundance in this model.^29^ This may have contributed to our inability to observe a transcriptionally distinct cluster of fusion competent pCT in bioreactor organoids or this could be due to rapid transit through the fusion competent pCT state. Time dependent sequencing of trophoblast organoids could yet reveal a coherent fusion competent cluster. Critically, consistent with other TSC-based organoid models, our pseudotime analyses suggest that CT27 and CT29 TSC organoid models may bypass a pre-fusion/early ST state (Sup. Fig 10).^28^ Using scRNA-seq data to model villous lineage trajectory, Shannon *et al.* observed that when TSC lines are used to produce Matrigel-based trophoblast organoids, TSCs may bypass pCT states and differentiate directly into ST.^28^ Interestingly, their data also revealed that primary cell derived Matrigel trophoblast organoids more faithfully represented *in vivo* villous lineage trajectory modelling. Therefore, data is consistently suggesting the widely available TSC-lines may use additional signaling pathways beyond those predicted using tissue and primary cells to form ST. This is congruent with our data showing *PRKCZ* KD did not impair ST fusion in the organoids despite reproducible effects on ST formation in explant cultures that rely on primary pCT. The difference in our results between models identify that a spectrum of pathways regulates this key process, and in some contexts the role of aPKC-ζ III in regulating fusion can be circumvented.

Interestingly, while *PRKCZ* KD decreased ST regeneration, inhibiting aPKC kinase activity did not impair explant and primary first trimester pCT fusion. APKC-ζ III likely only has basal levels of activity due to its inability to become activated via Par-6 mediated interactions. Graybill *et al.* created an aPKC mutant without a PB1 domain (only 2 amino acids smaller than aPKC-ζ III) and revealed the mutant lacked of kinase activity, supporting our hypothesis by suggesting that aPKC-ζ ΙΙΙ can participate in cell signaling by modulating binding interactions without the ability to phosphorylate targets.^19^ Importantly, the aPKC inhibitor used in this study blocks both aPKC-ζ and aPKC-ι activity, suggesting that both of these isoforms do not impact pCT fusion, or that they have opposing actions at this regulatory point.^54,55^ Since reintroduction of aPKC-ζ III alone in *PRKCZ* KO cells rescued fusion, it seems likely that the activity of both aPKC-ι and -ζ are not necessary at this point in ST formation, but further experiments are necessary to fully address the role of all aPKC isoforms and kinase-independent binding contributions. Our immunoprecipitation data revealed that aPKC-ζ III must be in excess abundance to LATS1 to significantly impair LATS1 interaction with Par-3. In the mouse brain, PKM-ζ (a brain specific *PRKCZ* encoded aPKC-ζ isoform) competes with aPKC-λ (aPKC- homolog) for binding to Par-3 to suppress axon specification, however, selective silencing of PKM-ζ allows for the maturation of a single axon.^56^ We suspect the upregulation of aPKC-ζ III is a critical step during villous lineage differentiation that is required to allow aPKC-ζ III to outcompete other cytoplasmic Par-3 binding partners to allow for Hippo signaling to occur.

SnRNA-seq data revealed an apparent stepwise increase in expression of *PRKCZ* mRNA from bipotential pCTs, pCTs, to fusion competent pCTs, suggesting there is an unknown regulatory stimuli controlling aPKC-ζ III expression. A stepwise increase in the expression of aPKC-ζ III was also seen in our TSC organoid model. Currently, it is not known if aPKC-ζ III encoding mRNA is transcribed from an alternative promoter sequence or via alternative splicing alone. In neurons, the expression of aPKC-ζ and PKM-ζ is epigenetically regulated via histone acetylation and DNA methylation.^57,58^ PKM-ζ is transcribed from an internal promoter sequence with a CREB binding site that is demethylated in differentiated neurons.^58^ Interestingly, CREB has been shown to play an important role in pCT differentiation, directly regulating *GCM1*, a key transcriptional regulator of ST formation.^59–61^ Therefore, understanding if aPKC-ζ ΙΙΙ is regulated via this alternative promoter, and therefore sensitive to cAMP/CREB activation will be important to examine in the future.^50^ The observed increased and sustained expression of *PRKCZ* mRNA and aPKC-ζ III protein in ST also highlights that this form of aPKC may have other critical roles in ST. We previously observed that antibodies raised against aPKC-ζ isoforms reveal a strong cytoplasmic ST signal and a cytoplasmic signal within first trimester pCT. ^24^ Together with the exclusive localization of aPKC-ζ III-GFP to the cytoplasm observed here, we can infer that aPKC-ζ III localizes to the cytoplasm in pCT and the ST.^24^ Without a polarized distribution, it is unlikely aPKC-ζ III directly regulates cell polarity in trophoblasts, though indirect regulation of polarity by competing for Par-3 binding may be possible.

Importantly, our work has shown a potentially trophoblast-specific regulatory point for the Hippo signaling pathway. Hippo signaling is a ubiquitous pathway and dysregulation has been well associated with numerous diseases such as immune dysfunction, cardiac disease, and cancer.^62–64^ APKCs have also been recognized as critical regulators of tumorigenesis and can have cell and cancer dependent tumor promoting or suppressive roles.^65^ Increased expression of aPKC-ζ has been found to promote breast cancer,^66^ colorectal cancer,^67^ and pancreatic cancer.^68^ Additionally, *PRKCZ* splice variants have been identified in prostate cancer.^69^ Reactivation of placental specific genes and pathways has become increasingly observed in cancer.^70,71^ Like the placenta, cancer cells have the ability to become invasive via epithelial-mesenchymal transition, induce tolerance of the immune system, and become multinucleated via the reactivation of genes encoding the syncytins, the retrovirally co-opted trophoblast fusogens.^70–72^ The potential for reactivation of aPKC-ζ III expression in pathology and the role it could play in modulating Hippo signaling in other tissues and cell types is another interesting future direction from this work.

Here we have identified, for the first time, a placental specific regulator of Hippo signaling. Dysregulation of Hippo signaling has been established in trophoblast dysfunction, resulting in placental pathologies.^73–77^ PE and IUGR are serious pregnancy disorders that complicate 2-8% of all pregnancies. While the etiology remains unclear, these complications are thought to originate from the placenta. ^78^ Trophoblast fusion and expression of fusion competency machinery are impaired in both PE and IUGR, thus, understanding the molecular regulators of trophoblast fusion is critical.^6,79,80^ Our work highlights the importance of discovering fundamental human biology and placental specific pathways for the development of therapies to treat placental pathologies. Thus, future studies determining how aPKC-ζ III expression is regulated and what additional roles it plays in villous trophoblasts may have widespread implications for human development and pathogenesis.

## Acknowledgments

We would like to thank the patients who donated tissue to our study and the staff of the Woman’s Health Options Clinic for the help in accessing this critical resource. We would also like to thank Dr. Mike Wong in the Advanced Cell Exploration Core for their support with the bulk RNA-sequencing and CRISPR-Cas 9 experiments.

M.R. is supported by the Canada Research Chairs program and support for equipment was provided to M.R. by the Canadian Foundation for Innovation. S.Z.S received salary support from Alberta Innovates and Advanced Education, and a Women and Children’s Health Research Institute Graduate studentships. W.K.D was supported by the Natural Sciences and Engineering Research Council of Canada and Women and Children’s Health Research Institute studentships.

Bulk RNA-seq library preparation was performed by the University of Alberta Faculty of Medicine & Dentistry High Content Analysis Core (RRID:SCR_019182), which receives financial support from the Faculty of Medicine & Dentistry, the Li Ka Shing Institute of Virology and Canada Foundation for Innovation (CFI) awards to contributing investigators.

Flow Cytometry Facility Experiments were performed at the University of Alberta Faculty of Medicine & Dentistry Flow Cytometry Facility, RRID:SCR_019195, which receives financial support from the Faculty of Medicine & Dentistry and Canada Foundation for Innovation (CFI) awards to contributing investigators.

Cell Imaging Core Experiments were performed at the University of Alberta Faculty of Medicine & Dentistry Cell Imaging Core, RRID:SCR_019200, which receives financial support from the Faculty of Medicine & Dentistry, the University Hospital Foundation, Striving for Pandemic Preparedness – The Alberta Research Consortium, and Canada Foundation for Innovation (CFI) awards to contributing investigators.

Plasmid sequencing was performed at the University of Alberta Faculty of Medicine & Dentistry Advanced Cell Exploration Core, RRID:SCR_019182, which receives financial support from the Faculty of Medicine & Dentistry, the Li Ka Shing Institute of Virology, Striving for Pandemic Preparedness – The Alberta Research Consortium, and Canada Foundation for Innovation (CFI) awards to contributing investigators.

## Author contributions

S.Z.S., W.K.D., and M.R. conceptualized the project. M.R. obtained funding for the project. S.Z.S., W.K.D., and I.D.K. performed experiments and analyzed data. J.G.R. performed all bioinformatic analyses. S.Z.S. and M.R. wrote the original manuscript with comments and edits from all authors. Supervision under M.R.

## Methods

### RESOURCE AVAILABILITY

#### Lead contact

The lead contact for this study is Meghan Riddell. All requests for reagents and methods should be directed to.

#### Materials availability

Plasmids generated from this study can be purchased from Vectorbuilder. *PRKCZ* KO lines can be requested from lead contact.

#### Data and code availability

Bulk-RNA seq and snRNA seq data generated from this study will be deposited and publicly available at the time of publication.

All original code will be published and available at the time of publication.

### EXPERIMENTAL MODEL AND STUDY PARTICIPANT DETAILS

#### Human placental tissue collection

Human placental tissue from weeks 5-12 gestational age was obtained from elective pregnancy terminations with informed patient consent according to methods approved by the University of Alberta Human Research Ethics Board (Pro00089293). All placental samples and patient characteristics for samples in this study can be found in Table 1. Biological sex was determined for 14 placental samples; 57.14% were male, 42.86% were female.

#### Bulk RNA sequencing

Libraries were prepared using the Illumina Stranded mRNA prep kit. Partial lane sequencing was performed by the Genome Science Centre (GSC) at the University of British Columbia via Illumina next generation sequencing (NovaSeq X Plus Series PE150) for 100M read pairs per sample. Sequencing data have been deposited in GEO and will be released upon publication.

Quality control and library alignment of the samples was performed in Ubuntu via the FastQC and STAR distros, respectively.^82^ We utilized R for the downstream analysis of the data. Counts were obtained utilizing the Rsubread package.^83^ Analysis of the data, estimation of fold changes and dispersion was performed using DESeq2. ^84^ The GO pathways were generated using the clusterprofiler package.^92^

#### Single nuclei sequencing

10x Genomics snRNA-seq techniques were used for flash frozen organoids, and we adapted the single nuclei sequencing data from first trimester placentas previously presented by Wang *et al.*^29^

Libraries were prepared by sequencing flash-frozen placental organoids by using a Chromium Nuclei Isolation with RNAse Inhibitor kit (Novogene, 1000494). Partial lane dual index sequencing was performed by the Princess Margaret Genomics Centre (PMGC) via Illumina next generation sequencing (Illumina NovaSeqX) for a total of 200M read pairs per sample. The sequenced data has been deposited in GEO and will be released upon publication. The first trimester single cell nuclei dataset was adapted from the sequencing data performed by Wang *et al.* in 2024.^29^

Sample demultiplexing, gene counting, and feature barcode analysis was performed on CellRanger-9.0.1.^89^ Downstream analysis was performed in R-4.5.1 and Seurat −5.3.0.^90^ Both datasets were filtered by excluding samples with less than 200 or more than 2500 features, as well as samples with more than 20% of mitochondrial counts. Dimension reduction was performed via UMAP, and the cell clusters were identified utilizing canonical markers for the cell identities.

Filtering was performed in both samples, and we retained a total of 45,697 nuclei for the first trimester dataset, and 22,250 nuclei for our organoid dataset. Batch correction and integration were performed in both datasets. The datasets were visualized using UMAP dimensional reduction analysis. For both datasets, clusters were annotated into the appropriate cell type by analyzing the expression of different canonical cell markers previously described (Figure S1-2).^30,36^

The pseudotime analysis was performed using the R package Monocle3.^44^ Trajectory reconstruction was performed in both the first trimester and the organoids dataset, with the bipotential pCTs population in the former and the pCT population in the latter used as the root for trajectory inference.

#### Explant syncytiotrophoblast regeneration

Human placental tissue was obtained with informed consent from individuals undergoing elective terminations. Tissue pieces (∼ 2mm^3^) were denuded of ST and cultured as per Duan *et al*.^36^ Briefly, placental tissue was washed with cold PBS cut into explants, then trypsinized (0.25% trypsin-EDTA, Gibco, 25200-056). Explants were cultured one/well, triplicate per treatment in explant regeneration medium [Iscove’s

Modified Dulbecco’s Medium (IMDM; Gibco, 12440061) supplemented with 10% FBS, 1% ITS-X (Gibco, 51500-056), and 50µg/mL gentamycin (Gibco, 15750-060)] in humidified incubators with 5% CO_2_ and atmospheric O_2_. After 24 hours, the tissue was vigorously washed to remove ST, debris, and treated with *PRKCZ* – targeting siRNA KD (Dharmacon, J-003526-14), non-targeting control siRNA (Dharmacon, D-001810-10), 5 µM myristoylated aPKC pseudosubstrate inhibitor (Invitrogen, 77749), or 3µM LATS inhibitor (TDI-011536; MedChemExpress, HY-150042). After an additional 48 hours of culture, explants were fixed with 4% PFA for immunofluorescence staining or collected for western blotting.

#### Generation of CRISPR-Cas9 *PRKCZ* Knockout BeWo Choriocarcinoma Lines

The Alt-R CRISPR-Cas9 System was used to create *PRKCZ* KO lines. *PRKCZ* targeting guide RNAs (CRISPR RNA; crRNA) were designed using the IDT Alt-R CRISPR HDR Design Tool to target base pairs 100584-100603 (tccagta gacgacaaga a) on the *PRKCZ* gene. Control lines were created using the Alt-2 Cas9 Neg Ctrl crRNA #1 (IDT, cat# 1072544, lot#0000828510). A two-part guide RNA, crRNA and Alt-R CRISPR Cas9 tracrRNA-ATTO 550 (tracrRNA; IDT, cat# 1075927, lot: 00005558022) were combined in equimolar concentrations for a final duplex concentration of 1µM in Nuclease-Free Duplex Buffer (IDT, cat#11-01-03-01). Ribonucleoprotein (RNP) complexes were transfected with Lipofectamine CRISPRMAX Transfection Reagent (Invitrogen, Cat# CMAX00001, lot 2634820) according to manufacturer’s protocols.

Briefly, Alt-R S.p. Cas9 nuclease V3 (IDT, cat#1081058, lot 0000833182) and crRNA/tracrRNA duplex were transfected using the Cas9 Plus Reagent (Invitrogen, Cat# 100035624, lot 2634802) and Lipofectamine CRISPRMAX Reagent (Cat# 100035629, lot 2650857) in Opti-MEM™ I Reduced Serum Medium (Gibco, 31985062). 24 hours later, cells were sorted via Fluorescence-Activated Cell Sorting Flow Cytometry for ATTO 550.

#### BeWo maintenance and *in vitro* syncytiotrophoblast differentiation

BeWo cells were maintained in Ham’s F-12 Nutrient Mix (Gibco, 11765047) supplemented with 15% fetal bovine serum (FBS; Wisent, Multicell Inc., Lot: 185730) and Penicillin-Streptomycin (100U/mL Penicillin, 100µg/mL Streptomycin; Gibco, 15140122) in humidified incubator at 37°C with 5% CO_2_ and atmospheric O_2_. To induce *in vitro* ST differentiation, BeWo cells were seeded on glass coverslips coated with 0.2% gelatin (Sigma, G1393). Medium was changed to also include 500µM 8-Br-cAMP (Sigma-Aldrich, B7880), and refreshed every 48 hours. Cells were fixed after a total of 96 hours of 8-Br-cAMP treatment. For western blotting assessment of p-YAP(Ser127) and p-LATS1(Ser909) expression, BeWo cells were seeded at 15% confluency, then transfected with aPKC-ζ III-FLAG the next day using Lipofectamine LTX reagent with PLUS reagent (Invitrogen, 15338100) in Opti-MEM I Reduced Serum Medium (Gibco, 31985062) according to manufacturer’s protocols. After 24 hours, cells were collected at 0 and 2 hours post 8-Br-cAMP treatment in RIPA supplemented with Protease Inhibitor Cocktail (Sigma-Aldrich, P2714) and Halt Phosphatase Inhibitor Cocktail (Thermoscientific, 78420).

#### BeWo siRNA knockdown

To perform *PARD3* knockdowns, BeWo cells were seeded on glass coverslips coated with 0.2% gelatin. After 24 hours, *PARD3-*targeting siRNA [ON-TARGETplus Human PARD3 (56288) siRNA; (Dharmacon J-015602-06)] and non-targeting controls, [ON-TARGETplus Non-targeting (Dharmacon, D-001810-10)] were transfected into BeWos using Opti-MEM™ I Reduced Serum Medium and Lipofectamine™ LTX Reagent with PLUS™ Reagent in Opti-MEM I Reduced Serum Medium according to manufacturer’s recommendations. The following day, *in vitro* ST differentiation was induced as above.

#### Human trophoblast stem cell culture

The human trophoblast stem cell lines^37^ (TSC; CT27, CT29) were obtained from Riken Biosource Resource Centre and maintained on 5µg/mL collagen IV (Corning, 354233) coated plates and cultured with human TSC culture medium ^36,37^ in a humidified incubator at 37°C with 5% CO_2_ and atmospheric O_2_. Human TSC lines were used from passage 20-29 and split at 1:5 – 1:20 ratios.

#### Human trophoblast organoid culture

TSC CT27 (Female line) and CT29 (Male line) cells were passaged and cultured in High Aspect Ratio Vessels (HARVs) (Synthecon) using a Rotary Cell Culture System (Synthecon). Organoids were pelleted via centrifugation, washed with PBS, and prepared for downstream analyses. For single nuclei RNA-sequencing, PBS was removed and the organoids were flash frozen. For western blotting, organoids were lysed in RIPA buffer. For immunofluorescent staining, organoids were fixed in 4% PFA for 10 minutes in siliconized 2mL round bottom tubes (Sigma, SL2).

#### *PRKCZ* knockdown organoids

*PRKCZ* – targeting siRNA KD [ON-TARGETplus Human PRKCZ (5590) siRNA (Dharmacon, J-003526-14)], or non-targeting controls, [ON-TARGETplus Non-targeting (Dharmacon, D-001810-10)] were transfected into TSC using Opti-MEM™ I Reduced Serum Medium and Lipofectamine™ LTX Reagent with PLUS™ Reagent according to manufacturer’s recommendations. 24h post transfection, treated TSCs were moved into HARVs for 24-48 hours in rotational culture.

#### Primary first trimester human progenitor cytotrophoblast culture and *in vitro* syncytiotrophoblast differentiation

First trimester pCTs were isolated and cultured as previously reported.^24,55,93^ For primary *in vitro* 72 hour ST differentiation, pCTs were cultured in IMDM supplemented with 10% FBS and penicillin–streptomycin in a 5% CO_2_ . Cells were seeded for 4 hours, then washed and treated with 10 µM 8-Br cAMP in IMDM + 10% FBS and pen-strep overnight. The following morning, medium was changed for 8-Br-cAMP removal, and the cells were cultured for an additional 48 hours before fixation.

#### Immunofluorescence staining

##### Placental tissue

Placental explants and tissue were simultaneously permeabilized and blocked using blocking buffer [5% Normal donkey serum, 0.5% triton-X 100, and 1:100 human IgG (Invitrogen, 02-7102)], then incubated overnight with primary antibodies (see table). The following day, tissue was washed and incubated with secondary antibodies and/or Phalloidin and Hoechst 33352. Tissue was mounted using imaging spacers and Fluoromount-G (SouthernBiotech, 0100-01).

##### Human trophoblast organoids and 2D cells

Human trophoblast organoids were stained as previously described.^36^ Briefly, organoids or cells (BeWo and primary *in vitro* differentiated ST) were permeabilized, blocked in blocking buffer (5% NDS, 0.01% Tween-20, and 1:100 human IgG), then incubated with primary antibodies overnight (Table 2). The next day, they were washed then incubated with secondary antibodies and/or Phalloidin and Hoechst 33352. Organoids were mounted using imaging spacers and Fluoromount-G. Cells cultured on coverslips were mounted with Fluoromount-G.

#### Image Capture and Analysis

Confocal microscopy was used to capture explant and organoid images. Three regions per explant or five organoids per treatment were captured using a Zeiss Plan Apochromat-20×/0.8 M27 or Zeiss Plan Apochromat-63×/1.4 M27 oil lens on a Zeiss LSM-700 confocal microscope. Z-stacks of placental explants(15-55µm) were captured at 20x (2.02µm step size) and 63x (1.2µm step size) (30-60µm). Z-stacks of organoids (20-30µm) were captured at 20x (2.02µm step size). XY-images were captured at 20x. Images were analysed using Volocity Imaging Software (Quorum Technologies, version 7.0.0). For 2D cell fusion assessment, triplicate images per treatment were captured at 10x magnification using an Olympus IX2-UCB immunofluorescent microscope equipped with a Roper Scientific camera and aa Sutter Instruments Lambda DG-4 fluorescent lamp and cellSens Dimensions imaging software.

#### Fusion assessment

##### Cells

Images were blinded for treatments and the number of nuclei incorporated into multinucleated E-cadherin positive clusters were counted and divided by the total number of nuclei using Image-J and Volocity Imaging Software.

##### Explants

ST regeneration of placental explants was quantified as previously described.^36^ Briefly, explants stained for E-cadherin (pCT marker), Phalloidin (F-actin), and nuclei were used to assess fusion. Single 20x XY-plane cross sectional images were assessed. Single nuclei surrounded by E-cadherin were considered pCT, and multiple nuclei surrounded by phalloidin and E-cadherin signal were considered ST. For each treatment, the area of ST/Area of pCT was normalized to donor-matched 24 hour control trypsinized samples, then to regenerated controls.

#### Western blotting

Protein was collected using RIPA (150 mM NaCl, 1% Triton-x 100, 0.1% SDS, 50 mM Tris, and 0.5% Sodium deoxycholate) supplemented with Protease Inhibitor Cocktail (Sigma-Aldrich, P2714) and Halt Phosphatase Inhibitor Cocktail (Thermoscientific, 78420) for phospho-specific antibody detection. SDS-PAGE was run using 10-20µg protein. Membranes were blocked with 0.3% skim milk powder and incubated overnight with primary antibodies (Table 2). The following day, membranes were washed then incubated with secondary antibodies. After incubation with phospho-specific antibodies, membranes were stripped for 4×30 minutes (7.5g glycine, 0.5g SDS, 5mL Tween-20, pH 2.2 w HCl), washed, blocked, and re-incubated with non-phosphorylated antibodies overnight. Total protein was determined using Fast Green stain (0.001% Fast Green FCF (w/v), 30% methanol, 7% acetic acid), then destained (30% methanol, 10% acetic acid). Membranes were imaged using the Licor Odyssey CLx and analyzed using Image Studio (V5.5)

#### RNA isolation and Reverse-transcriptase and polymerase chain reaction

Organoids were harvested and RNA was extracted using TRIzol-chloroform extraction and purified using PureLink RNA Mini Kit (Invitrogen). Reverse transcription was performed using iScript cDNA Synthesis Kit (BioRad) with 1000ng RNA, and cDNA was diluted 1:10 for all reactions. RT-PCR reactions were performed using SYBR Green Universal Master Mix (Applied Biosystems) on a QuantStudio 3 Real-Time PCR System (Thermo Fisher Scientific) and analyzed using QuantStudio Design & Analysis Software (Thermo Fisher Scientific). The 2^−ΔΔCT^ method was used to calculate relative change in mRNA expression using housekeeping gene *RNA18SN1.*^94,95^

*SRY* expression was used to determine the biological sex of placental samples. The amplicon was generated by performing PCR using PCR Supermix (Invitrogen) (amplicon = 169 bp) run on a C1000 Touch Thermal Cycler (Biorad, 1851148). Gel electrophoresis was performed to detect amplicons and visualized using RedSafe Nucleic Acid Staining Solution (Froggabio).

#### Plasmids

aPKC-ζ III-FLAG and aPKC-ζ III-EGFP plasmids were constructed and packaged by VectorBuilder (Table 4). Vector IDs can be used to retrieve detailed information about the vector on vectorbuilder.com. pEGFP-N1-Par3 plasmid was a gift from Dr. Masanori Nakayama. pcDNA3 Lats1 (Nigg HS189) (LATS1-Myc Tag) was a gift from Erich Nigg (Addgene plasmid # 41156 ; http://n2t.net/addgene:41156 ; RRID:Addgene_41156).^10^

#### Immunoprecipitations

For Par-3-EGFP and aPKC-ζ III-FLAG immunoprecipitations, HEK293T cells were seeded at 30% density and transfected the following day with 1:1 plasmid ratio using Lipofectamine™ LTX Reagent with PLUS™ Reagent according to manufacturer’s recommendations. Cells were lysed with ice cold lysis buffer [50 mM Tris–HCl pH 7.4, 1% IGEPAL, 150 mM NaCl, and 1:100 Protease inhibitor (P2714,Sigma-Aldrich, St.

Louis, MO, USA)], incubated for 15 minutes with end-over-end rotation at 4°C, and centrifuged for 15 minutes at 14,000 RCF. Protein assays were performed using Pierce BCA Assay Kit (Thermofisher, 23227). 50µL of Anti-FLAG® M2 Magnetic Beads (Sigma-Aldrich, M8823) were incubated for 4 hours at 4°C with 500µg protein lysate. Par-3-EGFP, LATS1-Myc, and aPKC-ζ III-FLAG immunoprecipitations HEK293T cells were seeded at 10% density and transfected the following day plasmids for Par-3-EGFP IPs with LATS1-Myc and aPKC-ζ III-FLAG. Cells were lysed with ice cold lysis buffer modified from Lv *et al*.^40^ [50 mM Tris–HCl pH 7.5, 0.3% IGEPAL, 150mM NaCl, 1mM EDTA-disodium, 1:100 Protease inhibitor, and 1:100 Phosphatase inhibitor (Halt™ Phosphatase Inhibitor Cocktail, Thermo Scientific, 78420), then incubated for 30 minutes with end-over-end rotation at 4°C and centrifuged for 30 minutes at 14,000 RCF. 25µL of GFP-Trap® Magnetic Particles M-270 (ChromoTek, gtd20; LOT; LM0000172) were incubated overnight at 4°C with 500µg protein lysate. Protein was eluted by boiling with 1X SDS buffer.

#### Live Cell Imaging

HEK293T cells were transfected with Par-3-EGFP or aPKC-ζ III-EGFP, and 24 hours later images were captured on a Zeiss Celldiscoverer 7 with an Axiocam 712 mono camera and Zeiss Plan-Apochromat 20x/0.7 autocorr lens.

#### Statistical analysis

All statistical analyses were performed in GraphPad PRISM (Version 10.4.2). For all tests, the threshold for significance is p>0.05. Statistical tests used for each experimental design are stated within figure legends. All graphs and representative images from different placental tissue donors are from at least three biological replicates with three technical replicates, or cell lines with at least three experimental replicates.

### ADDITIONAL RESOURCES

#### Data availability

SnRNA seq data can be found at https://riddell-lab.shinyapps.io/single_nuclei_placenta/.

