## Supplementary material for "aPKC-ζ III promotes trophoblast fusion by altering Par-3 interactions with Hippo Signaling Kinase LATS1": Table 1

**Table 1. Placental and Maternal age of samples.**

|  | Placental Characteristics | | Maternal Characteristics | |
| --- | --- | --- | --- | --- |
|  | Mean | S.D. | Mean | S.D. |
| Age (mean+/- S.D.) | 9.34473684 | 1.900170315 | 26.7575758 | 6.260143284 |
| Range (min, max) | (5, 12.7) |  | (18, 41) |  |
| Count | 38 | | 38 | |

*Biological sex was determined for 14 samples; 57.14% were male, 42.86% were female.*
