## Supplemental files for "aPKC-ζ III promotes trophoblast fusion by altering Par-3 interactions with Hippo Signaling Kinase LATS1"

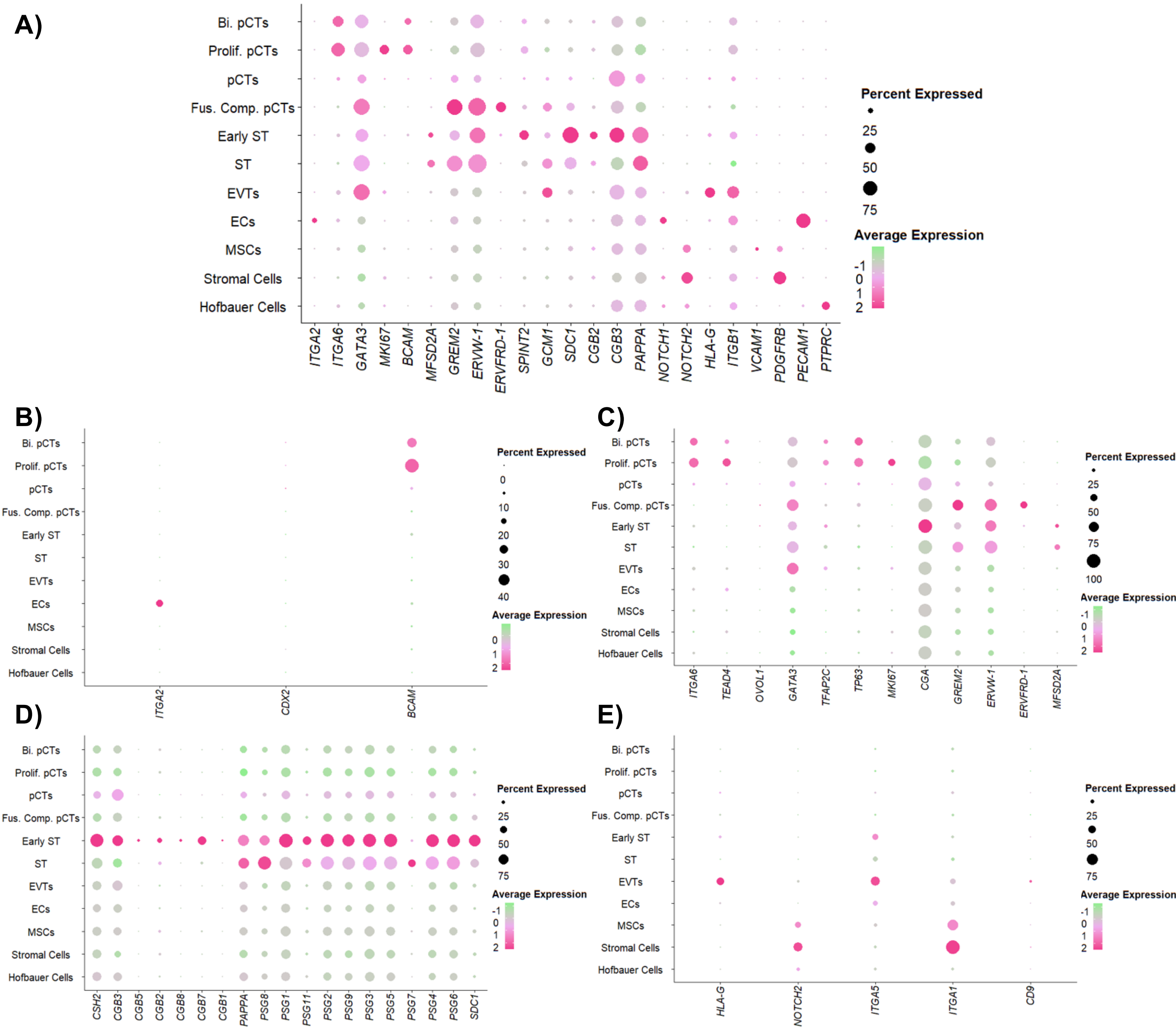


**Supplementary Figure 1: Subcluster defining marker gene expression in first trimester placenta.** Dot plots highlighting A) key marker genes, B) TSC, C) pCT, D) ST, and E) EVTs. PCTs = progenitor cytotrophoblasts; Bi. pCTs = bipotential pCTs; Prolif. pCTs = proliferative pCTs; Fus. Comp. pCTs = fusion competent pCTs; ST = syncytiotrophoblast; EVTs = extravillous trophoblast; ECs = endothelial cells; MSCs = mesenchymal stem cells.

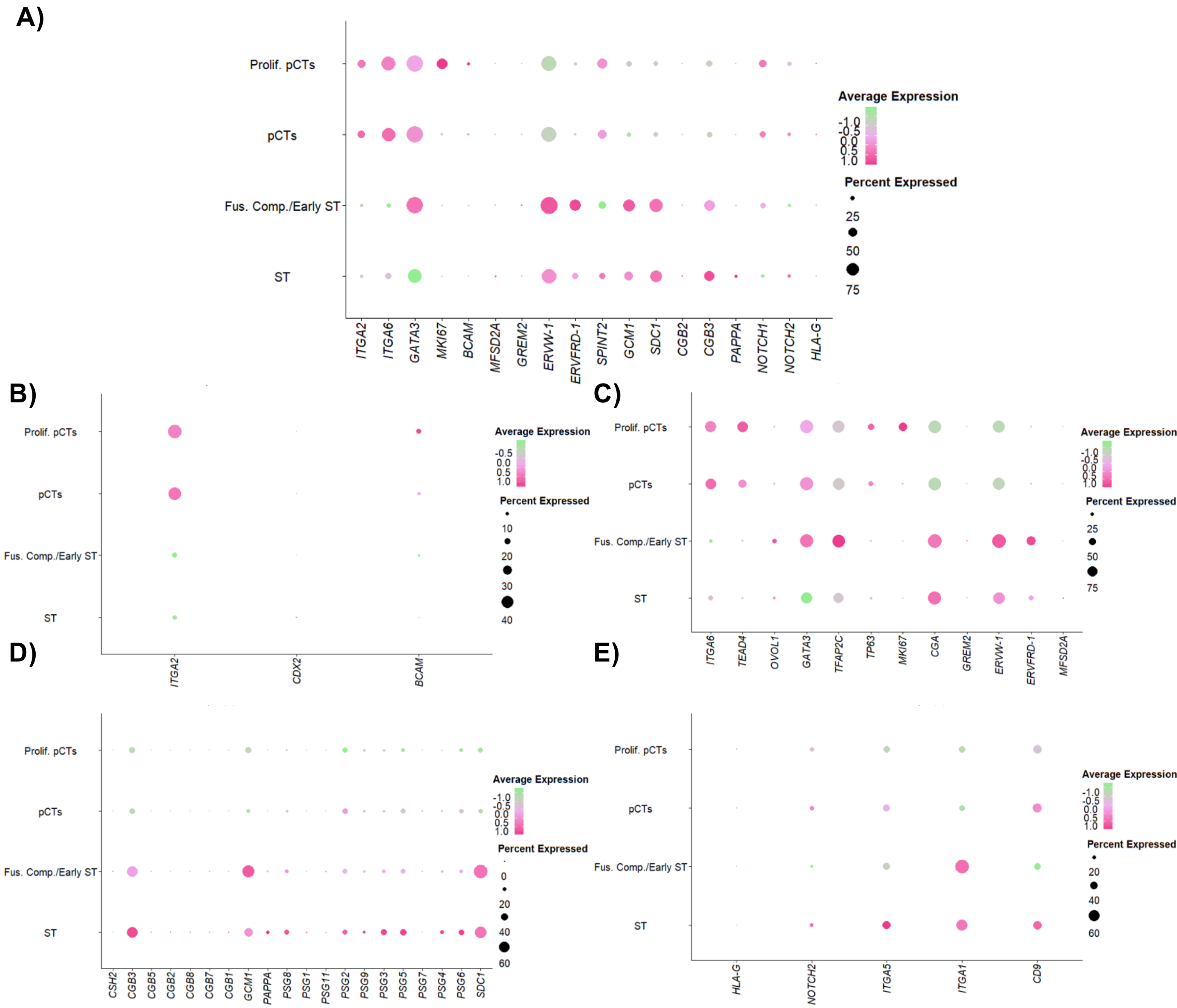


**Supplementary Figure 2: Subcluster defining marker gene expression in TSC organoids.** Dot plots highlighting key marker genes in A) all cell types, B) TSC, C) pCT, D) ST, and E) EVTs. PCTs = progenitor cytotrophoblasts; Prolif. pCTs = proliferative pCTs; ST = syncytiotrophoblast; Fus. Comp. pCTs / early ST= fusion competent pCTs / early ST.

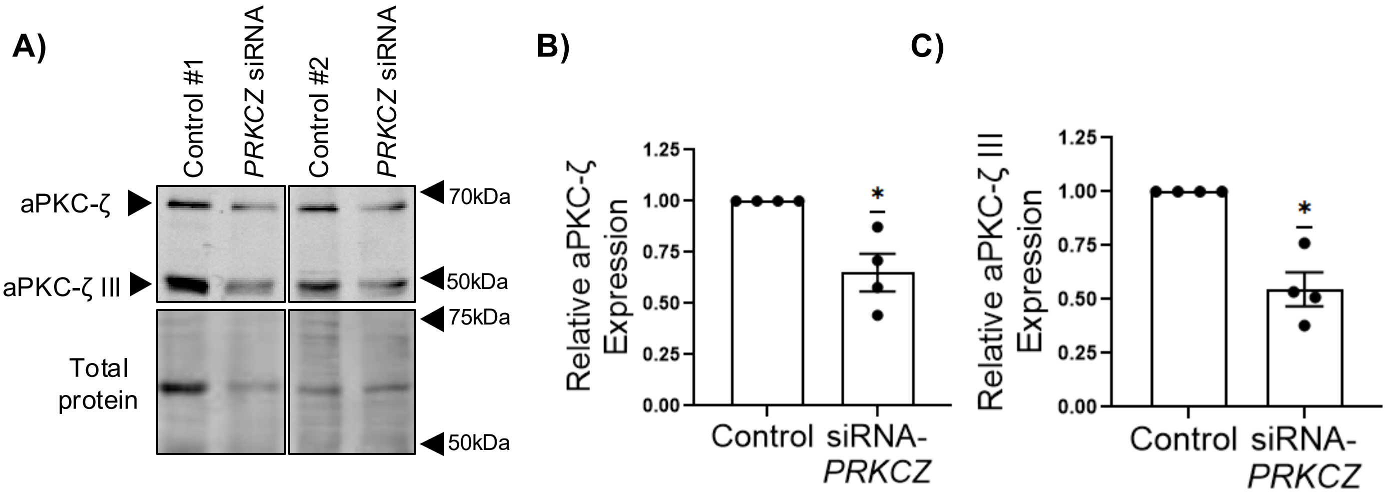


**Supplementary Figure 3: Knockdown of aPKC-ζ isoforms in placental explants** A) Representative western blot of aPKC-ζ and aPKC-ζ III in placental explants treated +/- *PRKCZ*-targeting siRNA; Summary data of relative B) aPKC-ζ andC) aPKC-ζ III expression; Data are mean +/- S.E.M., one sample t-test, *p ≤ 0.05, n=4.

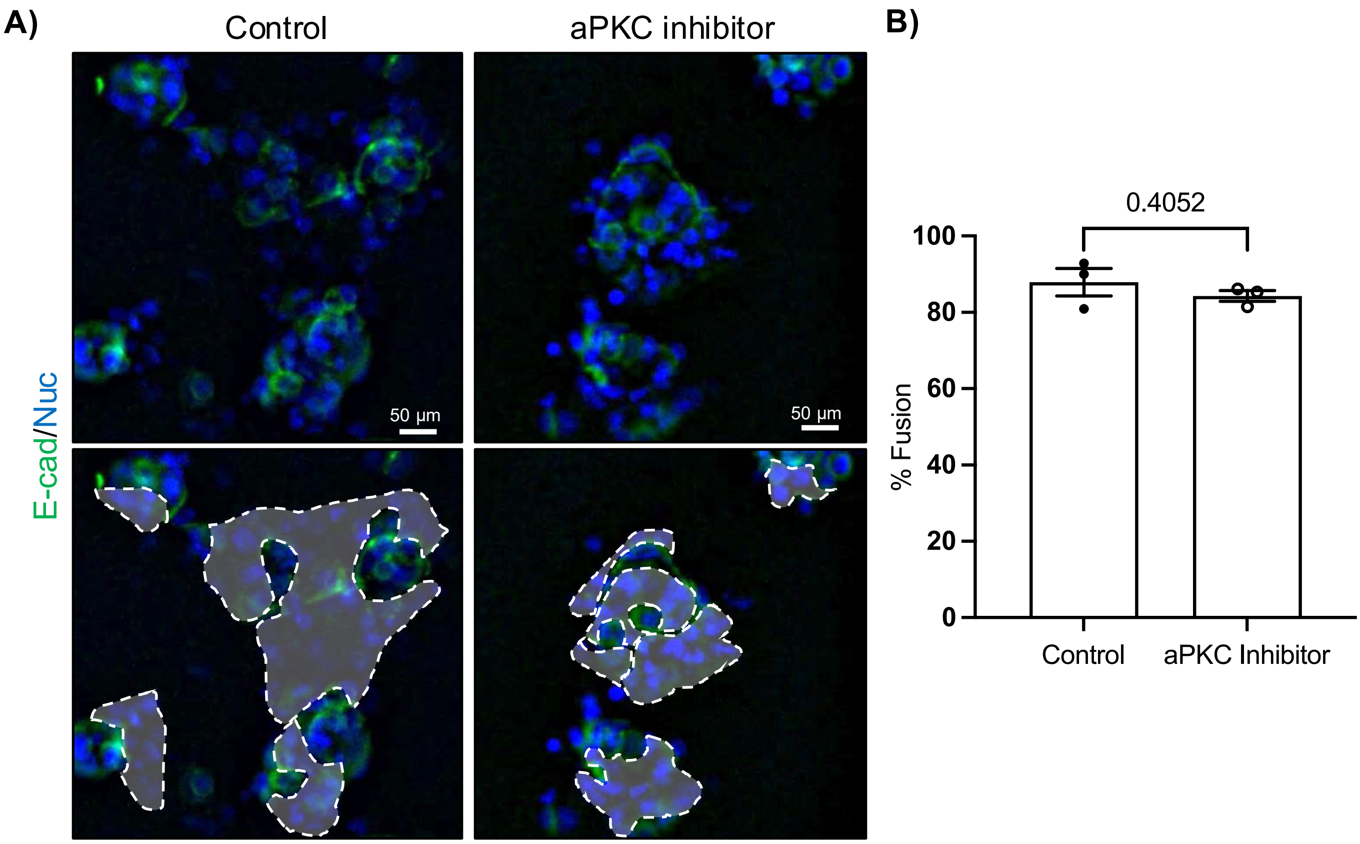


**Supplementary Figure 4: Trophoblast fusion in first trimester primary *in vitro* differentiated ST** A) Representative images of control and aPKC inhibitor treated primary first trimester in vitro differentiated ST stained for E-cad (E-cadherin; green) and nuclei; dashed regions below indicate regions of multinucleated cells. B) Summary data of percent fusion in control and aPKC inhibitor treated cells; Data are mean +/- S.E.M., paired t-test, n=3.

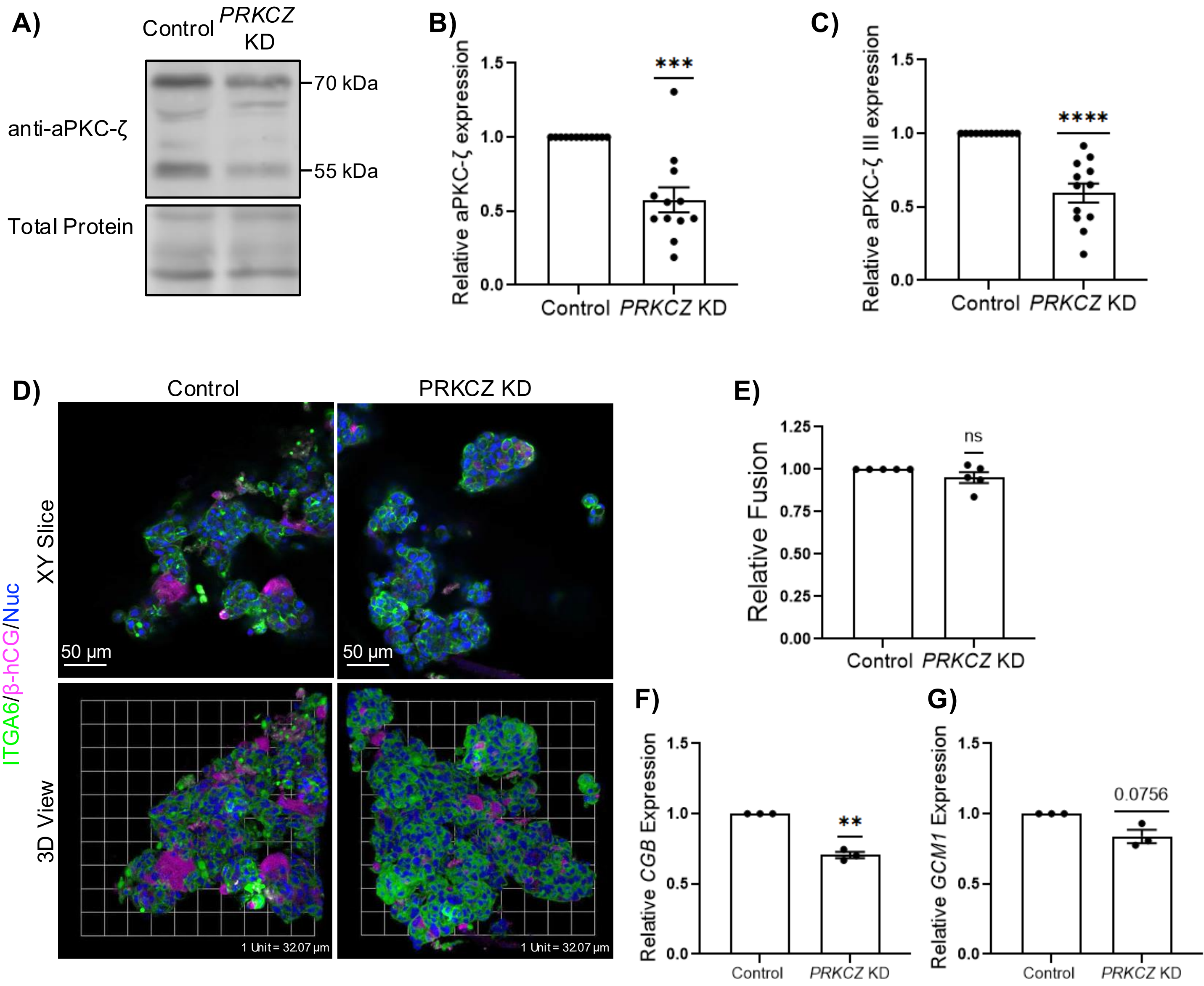


**Supplementary Figure 5: *PRKCZ KD* in human trophoblast organoids does not reduce fusion.** A) Representative western blot of aPKC-ζ expression in control and PRKCZ KD undifferentiated TSC. Summary data of relative B) aPKC-ζ and C) aPKC-ζ III protein expression; one sample t-test, n=11-12. D) Representative XY plane (top) and 3D reconstituted (bottom) confocal microscopy images of control and *PRKCZ* KD trophoblast organoids stained for ITGA6 (green), β-hCG (magenta), and nuclei (blue); scale bars = 50μm. E) Summary data for relative fusion of control and *PRKCZ* KD organoids;, one sample t-test, n=5. Relative expression of F) *CGB* and G) *GCM1* in control and *PRKCZ* KD organoids; one sample t-test, n=3. Graphs for all data are mean +/- S.E.M,; $^{*}p \leq 0.05$, $^{**}p \leq 0.01$ $^{***}p \leq 0.001$, $^{****}p \leq 0.0001$.

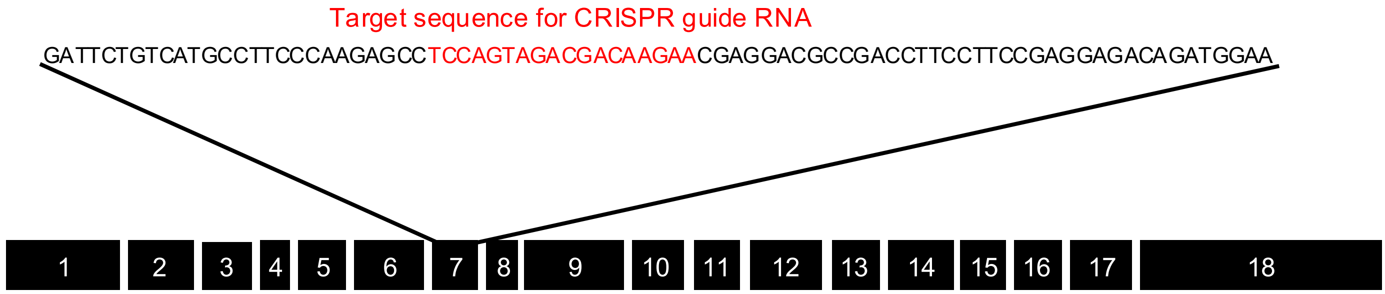


**Supplementary Figure 6: Schematic diagram of the *PRKCZ* gene with the 18 exons encoding for the full length aPKC-ζ.** Guide RNA target (Red highlighted sequence) for CRISPR Cas9 mediated knockout of *PRKCZ*. Exons 7-18 are conserved in aPKC-ζ III.

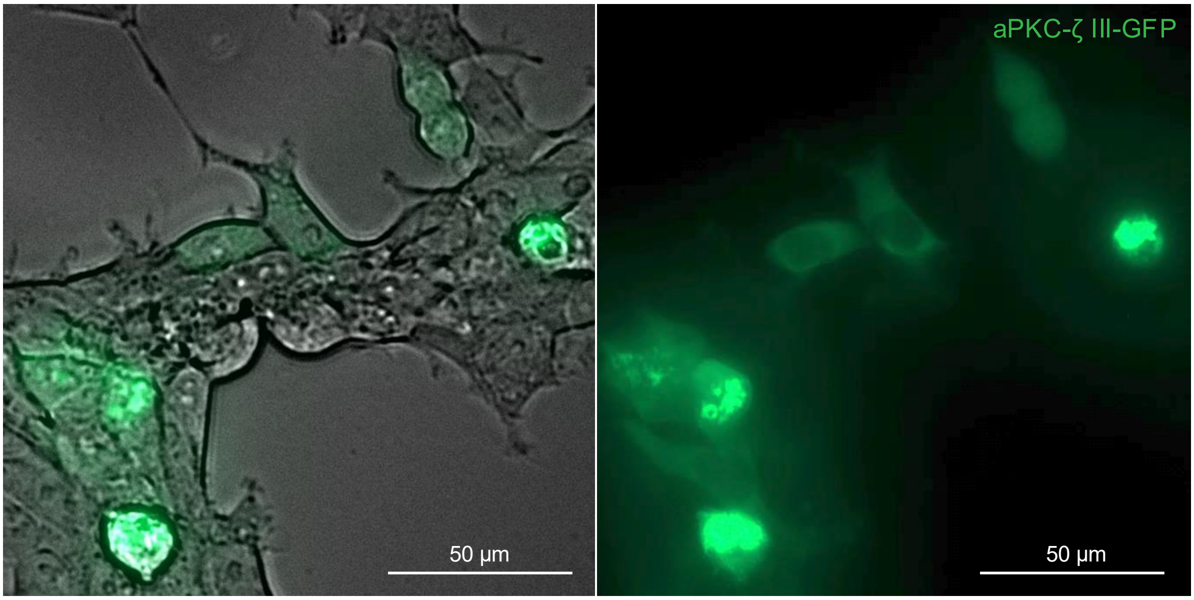


**Supplementary Figure 7: Live cell imaging of HEK293T cells transfected with aPKC-ζ III-EGFP plasmid.**

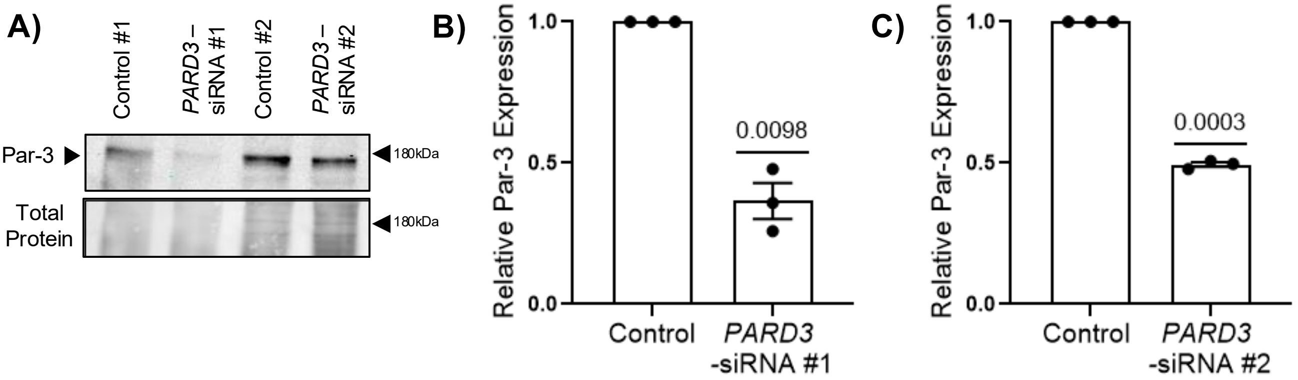


**Supplementary Figure 8: Par-3 KD in BeWo cells** A) Representative western blot of Par-3 expression in BeWo cells treated with *PARD3*-targeting siRNA; Summary data for relative Par-3 expression of B) *PARD3*-targeting siRNA #1 and C) *PARD3*-targeting siRNA #2; Data are mean +/- S.E.M., one sample t-test, n=3.

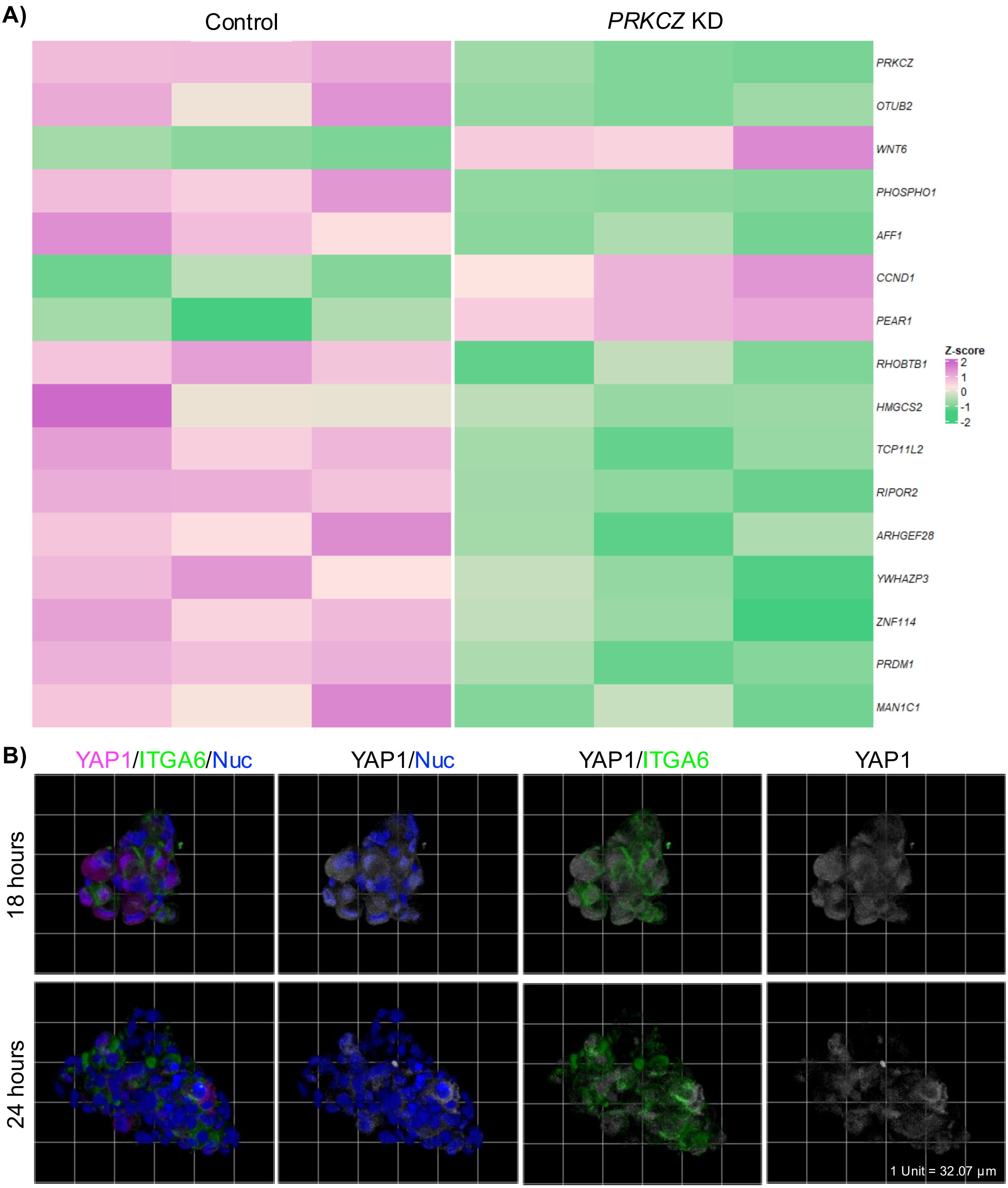


**Supplementary Figure 9:** A) Heatmap of significant differentially expressed genes in control and *PRKCZ* KD trophoblast organoids identified by bulk-RNA seq.. B) Representative 3D reconstituted confocal microscopy images of 18 and 24 hour trophoblast organoids stained for YAP1 (magenta), ITGA6 (green), and nuclei (blue).

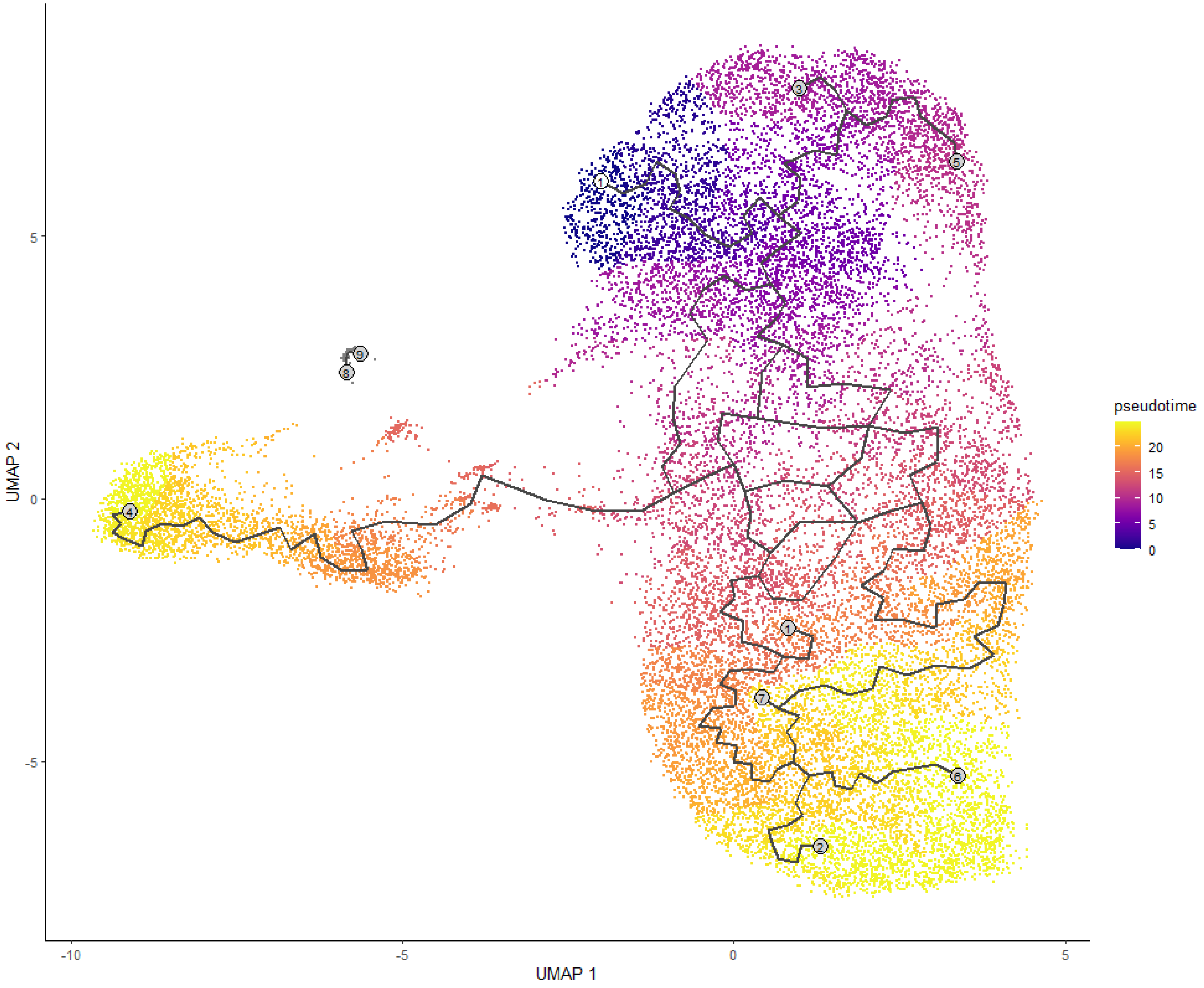


**Supplementary Figure 10: Pseudotime trajectory analysis of trophoblast organoids.**
